# A Vast Navigable Protein Sequence Subspace

**DOI:** 10.64898/2026.10.18.759989

**Authors:** Sam Sinai, Andreea Gane, Nina Jain, Drew Bryant, Max Bileschi, Ali Bashir, George M. Church, Eric Kelsic, Lucy Colwell

## Abstract

Fitness landscapes govern evolution. Our empirical understanding of their structure has primarily been informed by evolutionary history. Here, we combine high-throughput DNA synthesis and machine learning, without training on evolutionary priors, to build a high-resolution map of the genome packaging landscape of the Adeno-associated Virus capsid protein. We assay 232,123 mutants around Adeno-Associated Virus 2 (AAV2) including 15 neighborhoods centered on seed sequences within 5–21 mutations of the wild type sequence. Local neighborhoods differ sharply in mutational tolerance and epistatic detail, yet much of the quantitative variation is captured by a simple pattern: sitewise effects transformed by global epistasis. Strikingly, using targeted sampling of only a few thousand mutants, we uncover a vast percolating network of function-preserving ridges embedded within a largely non-functional landscape. These ridges support billions of distinct trajectories of up to 667 steps that connect all sampled neighborhoods through functional single mutations. Designed variants with mutations distinct from the evolutionary background retain packaging at distances where randomly generated mutants rarely survive. These include capsids that combine up to 7 mutations that are largely absent from the evolutionary record, and which are each individually lethal in the 15 diverse backgrounds we tested. Our results reveal rich subspace structure resembling *landscape orebodies*: reservoirs of highly viable and navigable sequence space defined by simple constraints. These subspaces can be discovered through targeted sampling and can open access to vast troves of proteins that may be valuable for downstream applications such as therapeutics.

## Introduction

Fitness landscapes map genetic sequences to phenotypic traits. Conceptualized nearly a century ago (Wright 1932) they can be seen as the solution space within which evolutionary search finds useful functionality. As such, understanding the structure of fitness landscapes has wide implications for evolutionary biology, population genetics, and increasingly synthetic biology and generative design of therapeutics (Bryant et al. 2021; Ogden et al. 2019; Bank et al. 2016; Sarkisyan et al. 2016; Gonzalez Somermeyer et al. 2022; Biswas et al. 2021; Zhou et al. 2022; Puszkarska et al. 2024; Papkou et al. 2023; Bakerlee et al. 2022; D. Ding et al. 2022). For most of their conceptual history, their study was underpinned by simple theoretical models, such as Random, Additive, and NK models (Kauffman and Weinberger 1989).

Recent treatments rooted in theoretical and empirical insights have focused on decomposing epistatic interaction based on order (Kauffman and Weinberger 1989; Otwinowski et al. 2018; Zhou and McCandlish 2020; D. H. Brookes et al. 2022; Reddy and Desai 2021; Poelwijk et al. 2019; Weinreich et al. 2013; Sailer and Harms 2017), and inferring empirical fitness functions to arbitrary accuracy, but often require dense sampling within a restricted slice of the mutational landscape. Rapid development in our ability to assay the functionality of many sequences provides a powerful tool to empirically investigate properties of fitness landscapes (Zhou et al. 2022; Sarkisyan et al. 2016; Bank et al. 2016; D. Ding et al. 2022; Bakerlee et al. 2022; Ogden et al. 2019; Gonzalez Somermeyer et al. 2022; Papkou et al. 2023). Yet, fitness landscapes, especially far from the sequence space explored by evolution, remain hard to interpret and understand; to make progress we must find a balance between the interpretability of simple models and the precise prediction ability and high flexibility that emerges from training high-capacity models on large datasets (Puszkarska et al. 2024; Hopf et al. 2017; Hsu et al. 2022; Madani et al. 2023; D. Ding et al. 2024; Candido et al. 2026; Park et al. 2024).

Several studies of this kind have emerged in the last decade, marrying high-throughput synthesis with high-capacity models (Bryant et al. 2021; Gelman et al. 2021; Romero et al. 2013; Wu et al. 2019; Hsu et al. 2022). Many are in the domain of protein design, where method development is a priority (Bryant et al. 2021; Puszkarska et al. 2024; Biswas et al. 2021; Zhu et al. 2024), but some also address more fundamental questions about fitness landscapes (Otwinowski et al. 2018; Papkou et al. 2023). Multiple studies explore mutants around GFP variants using systematic mutagenesis and modeling (Sarkisyan et al. 2016; Poelwijk et al. 2019; Gonzalez Somermeyer et al. 2022). Rotrattanadumrong and Yokobayashi explore an RNA fitness landscape and report a large neutral network connecting two functional mutants separated by 16 mutations (Rotrattanadumrong and Yokobayashi 2022). Due to their therapeutic importance, a series of recent studies have focused on validating and designing AAV capsid sequences with supervised and unsupervised models that can maintain productivity in high-throughput experiments (Ogden et al. 2019; Bryant et al. 2021; Zhu et al. 2024; Sinai et al. 2021). These studies showed that computational models can explore deep into the sequence space while maintaining capsid functionality. However, the objective of these studies was to develop methods and design proteins with high diversity and not to closely scrutinize the fundamental structure of the landscape.

Here, we characterize the local and long-range structure of a viral productivity fitness landscape (Fig. 1A) as an essential property needed for any downstream function (Adachi et al. 2014; Ogden et al. 2019). In other words, this landscape is the bedrock on top of which any secondary, and frequency-dependent traits would evolve. It is a functionally complex filter an AAV needs to pass at minimum to be viable in the wild or useful in therapeutics. We study the 28-residue region of the AAV2 VP capsid protein from positions 561 to 588, a region that includes both buried and surface-exposed residues and is of high relevance for in vivo behavior (Fig. 1B). The strong reproducibility of our assay (Fig. S1) allows us to measure quantitative differences in packaging efficiency among variants, increasing the resolution at which we can understand landscape structure. Interrogating the topography of a fitness landscape at a high edit distance can be challenging as a vast majority of sequences are completely non-functional - e.g. a majority of random single mutations on the AAV capsid break either protein folding, capsid assembly or genome packaging. However, generating functional sequences at high edit distance with any generative process also introduces bias because a specific model of the fitness space is needed to increase the viability of sampled variants. To navigate this trade-off, we base our investigation in this study on the neighborhoods surrounding 15 pre-selected, viable, sequence-diverse *seeds, distinct from natural variants.* These are variants that produce similarly or better than the wild type (WT), measured in (Ogden et al. 2019; Bryant et al. 2021) and possess significant diversity at the sequence level, with the farthest being 21 mutations away from WT AAV2. We explored these neighborhoods by (1) Stochastic (unbiased) sampling of mutants up to distance 5 to each seed, including all single substitutions and insertions, (2) Interpolating between seeds by taking direct combinatorial (stochastic) paths in sequence space from one seed to another, and (3) Moving away from each seed and multiple other starting positions in the landscape using machine learning models trained on the original data, using the same models as in (Bryant et al. 2021) (Fig. 1C).

**Figure 1.**
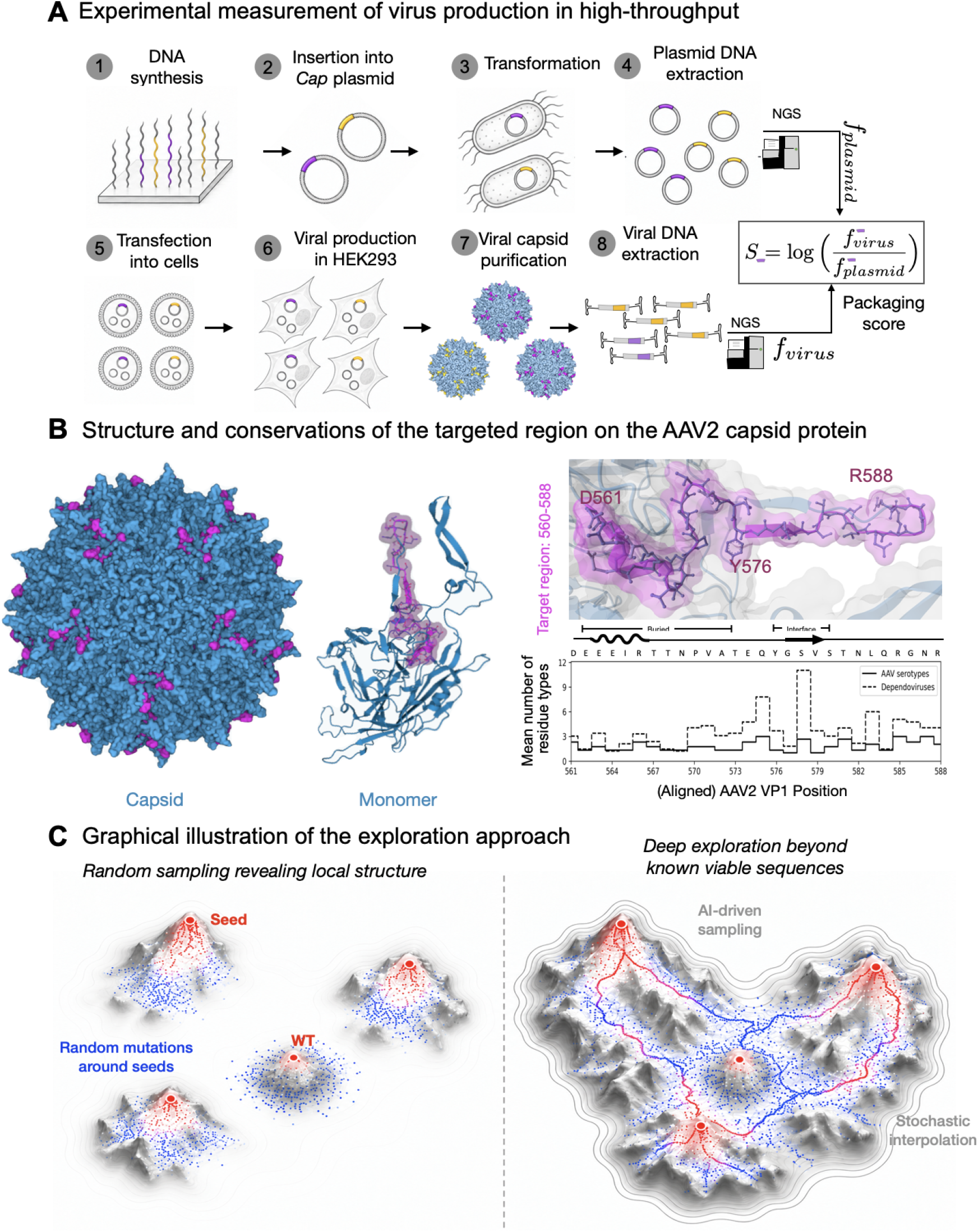
Overview of experimental design. A: A graphical summary of the measurement assay. Chip-based synthesized DNA is inserted into a Cap plasmid, and then transformed into *E. Coli*, cloned and extracted. Next-generation sequencing (NGS) determines the frequency of each variant within the plasmid pool. Plasmids are then transfected into HEK293 cells to produce viral particles. DNA from viral particles is then extracted and sequenced to calculate the frequency of variants in the virus pool. The packaging score (fitness) is the ratio between these measurements. B. The 28-residue target region contains both surface-exposed and buried residues relevant to viral production and function and therefore of high interest (Adachi et al. 2014; Ogden et al. 2019; Bryant et al. 2021). Mean number of residue types (perplexity) per position within the AAV serotypes and dependoparvovirus family. C: Data collected in each neighborhood contains every single substitution and single insertion, random variants at edit distance 2-5. In tandem we explore deep into the sequence space using stochastic, step-wise recombinant paths between seeds, together with machine learning driven exploration.

By combining local mutational sampling, stochastic interpolation between distant functional variants, and machine-learning–guided exploration, we uncover an unprecedented level of detail about a protein fitness landscape that is simultaneously rugged at short distances but highly smooth and navigable nonetheless. While individual mutational neighborhoods differ substantially in robustness and epistatic structure, much of the quantitative variation in fitness is captured by simple site-wise effects modulated by global epistasis. At the same time, direct step-wise paths between distant functional variants reveal an unexpectedly large space of functional connectivity in this protein landscape. Finally, we observe that ML models can find mutants that are categorically different from those likely to be discovered by evolution, and show that the landscape has structured subspaces with a relatively high density of functional variants, due to simple inclusion/exclusion rules. We use *landscape orebodies* as a metaphor to describe these subspaces and suggest that while the evolutionary record can provide a few initial points to start the sampling, the core structure of these regions of the fitness landscape are often absent from the evolutionary record, but can be learned efficiently through a few thousand samples.

## Results

We first characterize the landscape through the lens of random mutants around each seed sequence. We measure the neighborhood around each seed by sampling 1092 (all) single mutants, 3600 random double mutants, and 1633 random mutants at edit distances 3 to 5 to each seed (substitutions and insertions, 6325 mutants per seed in total). The closest seed is 5 mutations away from WT, and the farthest is 21 (Fig 2A).

**Figure 2.**
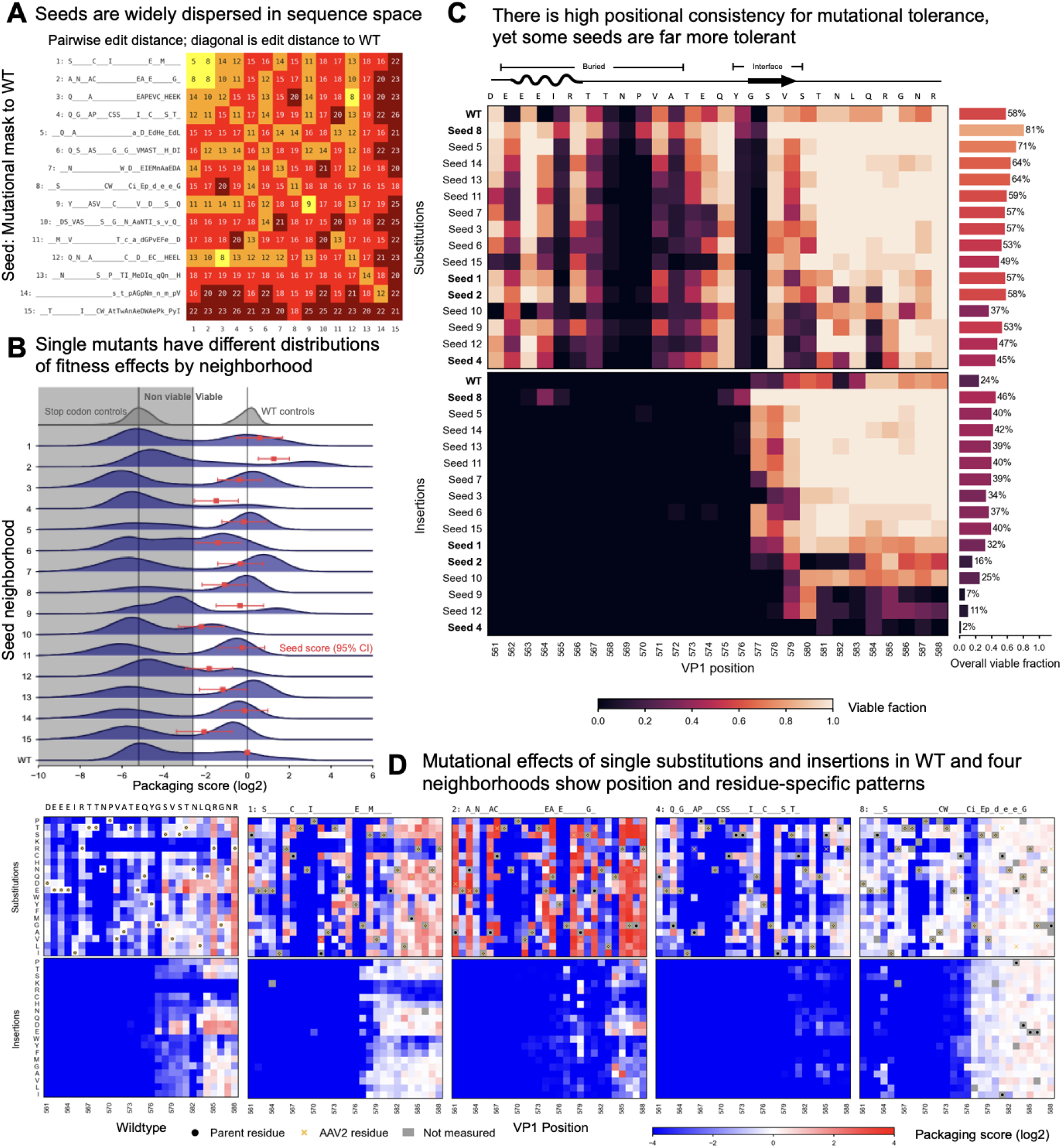
Mutational neighborhoods of different background sequences show shared fundamentals but diverse characteristics. A: Seeds span a large space within the 28-amino acid window, at 5-21 mutations (substitutions and insertions) to wild type and 8-26 mutations from each other. B: Distribution of fitness effects of all single mutations around each seed, compared to controls (WT replicates (n=200) and stop codon replicates (n=171)). C: Proportion of viable variants for single-mutant substitutions and insertions per position. Seeds are sorted by their total viability (substitutions and insertions). D: Mutational effects of single substitution and insertions per residue for WT and four different seeds (See Fig. S2 for all 15)

### Local neighborhoods reveal shared patterns and unique characteristics

Examining single mutations around each seed (Figure 2) certain fundamental patterns emerge (see Fig. S2 for a complete map of single mutants for all seeds). Specifically, the second half of the region demonstrates greater tolerance for substitutions and insertions compared to the first 15 positions. This observation aligns with prior findings (Bryant et al., 2021) and is consistent with the structural context, as most residues in the first 15 positions are buried (Fig. 1B). Additionally, the mean number of residue types, which reflects positional conservation (also referred to as perplexity), is lower in the first half of the region (Fig. 1B).

Despite these shared characteristics, significant diversity is also evident across seeds. For instance, seeds with similar fitness levels exhibit distinct mutational responses: seed 8 demonstrates high robustness to single mutations while seed 4 is comparatively brittle (Fig. 2B). Substitutions at residue 561 of seed 2 generate exceptionally high scoring mutants, while most of the same substitutions for seed 10 will result in non-viable capsids.

Variation in tolerance to insertions further highlights neighborhood-specific differences (Fig. 2C). Insertions are highly deleterious for seeds 2, 4, 9, and 12, even though substitutions are tolerated. In contrast, seeds 3, 5, 7, and 14 accommodate or even benefit from single insertions, particularly in the last third of the target region. The impact of individual mutations also varies significantly across backgrounds. For example, mutations that are beneficial in the seed 7 context are neutral in seed 14 and deleterious in seed 10, despite these seeds having relatively similar fitness levels (Fig. 2C).

Certain amino acids also show distinct effects. Seed 1, for example, tolerates most insertions within the same region except for specific residues such as R, K, and W, which are tolerated in many other backgrounds. For instance, seed 8, carrying two negatively charged insertions on its backbone, tolerates these mutations well. Similarly, C is generally deleterious across most backgrounds but is notably tolerated by seed 3, with some tolerance also observed in seeds 7 and 8. Overall, for insertions and substitutions, position is far more explanatory of mutational effect compared to background- or residue-specific effects (Fig. S3 provides additional evidence).

Single mutations in all backgrounds can be roughly grouped into three classes: those that are generally tolerated, those that are tolerated in specific backgrounds, and those that cause a loss of function in every background (Fig S4.). Between 37.1% (seed 8, 11 mutations from WT) and 77.1% (seed 4, also 11 mutations from WT) of mutations break the packaging completely. These universally lethal mutations are highly informative in understanding the global structure of the landscape and are discussed later in the manuscript.

### Neighborhoods are described well by their own single mutant effects

The fitness effects of multi-mutants sampled randomly around each seed expose regularity in the local structure of the landscape. Across the 15 neighborhoods we measure 91,876 proteins, including all single mutants, and random 2-5 mutants near each seed (For the purpose of this analysis we reserve 15,443 measured multi-mutants as a hold-out). We observe that in all neighborhoods, viability drops quickly as more mutations are added and the independent-sites (Additive) models are highly predictive.

We first build descriptive models as tools to understand the landscape.

● **Single-hit model.** Assumes that it takes a single lethal mutation to break the capsid. In other words, the average probability *P_d_* of a viable random variant at edit distance *d* from seed *i*, can be calculated as a simple multiplication *P_d_ (viable; i)* = *P_1_ (viable; i) ^d^*. Note that viability is calculated within the seed background, rather than aggregated across seeds.
● **Sum of (measured) single-mutant effects.** In the basic case, simply adding what was measured (see below and Methods). It can be augmented by adding a simple non-linearity (e.g. a sigmoid) to account for global epistasis (Otwinowski et al. 2018).

The single-hit model explains a majority of the variation in the proportion of viables, suggesting that in most cases a single mutation, as opposed to negative epistasis, is sufficient to break the capsid. Pooled across neighborhoods, the viable fraction falls from 0.44 at one mutation to 0.016 at five, and the multiplicative prediction from the pooled single-mutant viability alone tracks it well (Fig. 3A).

**Figure 3.**
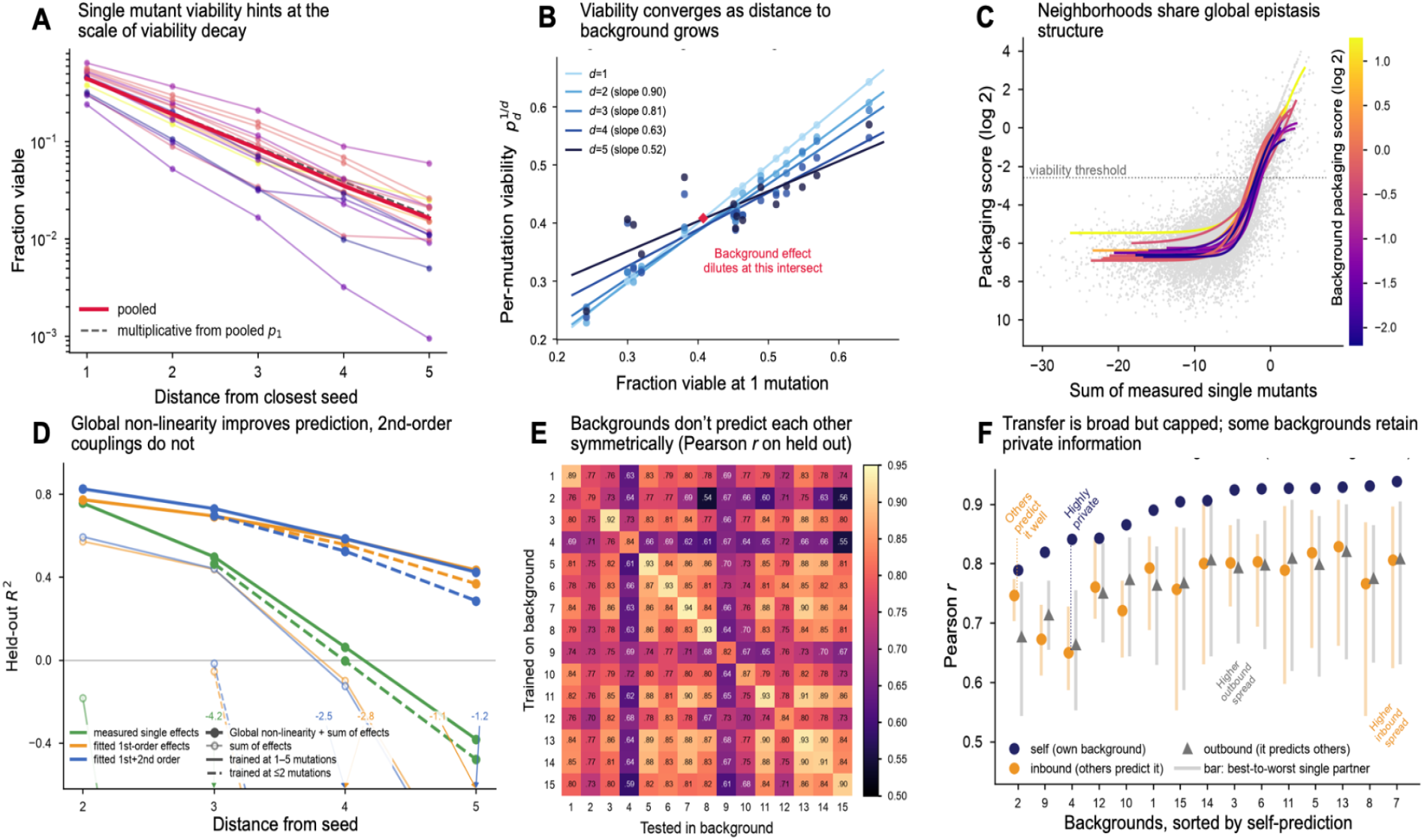
Multi-mutants are highly predictable from local single-mutant measurements, but most backgrounds keep private information. A: The fraction of single mutants viable *p_1_* around each seed is a helpful predictor of multi-mutant (at distance *d*) viability on average through *p ^d^*. B: Per-mutation viability *p ^1/d^* against the same background’s viability at one mutation. As distance grows background identity dilutes. C: Grey points: sum of the actual measured single-mutant effects against measured score for all variants; Curves are simple sigmoids fitted independently within each background and coloured by its packaging score. D: The sum of *measured* singles is strong at ranking multi-mutant performance (median *r* = 0.82), but quantitative prediction improves substantially through the simple non-linear transform (i.e. global epistasis). We find accounting for 2nd order effects to be of negligible importance in this dataset. E: Models trained in each background and scored on the target column’s held-out test proteins (diagonal and off-diagonal entries are comparable). While many pairs of backgrounds are symmetric in their predictive power, some small and some large asymmetries in inbound and outbound prediction are observed (e.g. (2, 8), (2,15)). F: In aggregate, inbound and outbound predictions on each model can also be distinct, and the gap between self-prediction and prediction from others can be large. Some backgrounds (e.g. seed 4) contain “private” information, not learned in other backgrounds and not useful to predict them.

While interpretive, this model is not perfect. If we regress each neighborhood’s per-mutation viability *p_d_ ^1/d^*_d_ on its viability at one mutation (Fig. 3B), the slope declines steadily with distance (i.e. drops from 0.9 at two mutations, to 0.52 at five, compared to a purely multiplicative model’s expected slope of 1). The fits anchor around a common per-mutation viability near 0.41: tolerant neighborhoods lose viability faster for each mutation than their own single-mutant rate would predict, and intolerant ones decay more slowly, so neighborhoods become progressively more similar as mutations accumulate. Fitting the decay of these slopes gives the identity of the background a half-life of ∼5 mutations (95% CI 2.5-10.6). In other words, single-mutant measurements set the local rate of viability decay, but that information is diluted as we move away from known seeds.

The sum of measured single-mutant effects moves us beyond viability alone. It tracks multi-mutant fitness (but does so non-linearly, Fig. 3C). Fitting a simple sigmoid non-linearity independently within each neighborhood recovers curves that are close to a shared one. Neighborhoods occupy different positions on a common non-linearity, which is what a generic model of a complex trait predicts (Reddy and Desai 2021). The single-hit and sum-of-singles models converge under certain circumstances, such as when a single mutation alone is so deleterious that beneficial mutations fail to rescue the variant.

Per neighborhood, the global epistasis model built from measured singles explains held-out double mutants with R2 >= 0.7 in 11 of 15 cases, while the additive score alone ranks the same variants well (median r = 0.82) but is quantitatively off in most neighborhoods. As more mutations are introduced the explanatory power of all these models drops, though this is also a consequence of most variants being non-functional, which makes measurements noisier. We also test for pairwise interactions among newly introduced mutations, and find them to be negligible in determining fitness (Fig. S5).

We then ask how much information is shared among neighborhoods by training a global epistasis model with fitted first-order effects in one neighborhood and evaluating it in every other (Fig. 3E). The neighborhoods share substantial structure. The mean cross-background correlation is 0.768, against 0.891 for models trained and tested in the same neighborhood, and the worst pair in the matrix still reaches r = 0.54. We hold out the same test set (∼1000 variants) for each neighborhood, enabling direct comparison between self and other predictions. While the matrix is close to symmetric, the direction of transfer can matter for individual pairs. 23 of the 105 neighborhood pairs do differ significantly in the two directions, and 9% of the variance of the matrix, well above the noise threshold, is due to this asymmetry.

All but one neighborhood stores some “private” information: information that is not predicted, or predictive of other neighborhoods (Fig. 3F). Self-prediction beats the average of the other 14 neighborhoods in all 15 cases (mean margin 0.123, Wilcoxon p = 3.1 x 10^-5) and the single best source (mean margin 0.049). Neighborhood 12 is an exception where there is a near-tie against the single best source. Across neighborhoods, self-prediction quality follows single-mutant viability (r = 0.78) but not the training-set size (r = -0.34). While generally inbound and outbound quality are strongly coupled (r = 0.838), there are exceptions. Neighborhood 2 shows significant differences between inbound and outbound, where it’s easier to predict it compared to how it predicts others. Neighborhood 4 hides the most information from others, and neighborhood 3 has a higher outbound spread than inbound, whereas neighborhood 8 is the opposite.

The private information is not from pairwise effects from newly introduced mutations (Fig. S5a) or how well the background is explained by global epistasis (Fig. S5b), but rather the idiosyncratic effects of single mutations given the background they are set in, which cannot be learned from the set of mutations introduced by random sampling alone.

### A viability-preserving network connects every seed

While random exploration away from a seed collapses quickly, interpolation between distant functional seeds reveals the most unexpected property of the landscape. In this section, we inspect the landscape through stochastic pairwise paths between seeds, where at each step we swap each mutation in one source sequence with the mutation in the target seed (Fig 4A). We sample these paths both in dense mode (4,993 samples, every single-mutant step is included) and sparse modes (22,500, only some hops along the path are included).

**Figure 4:**
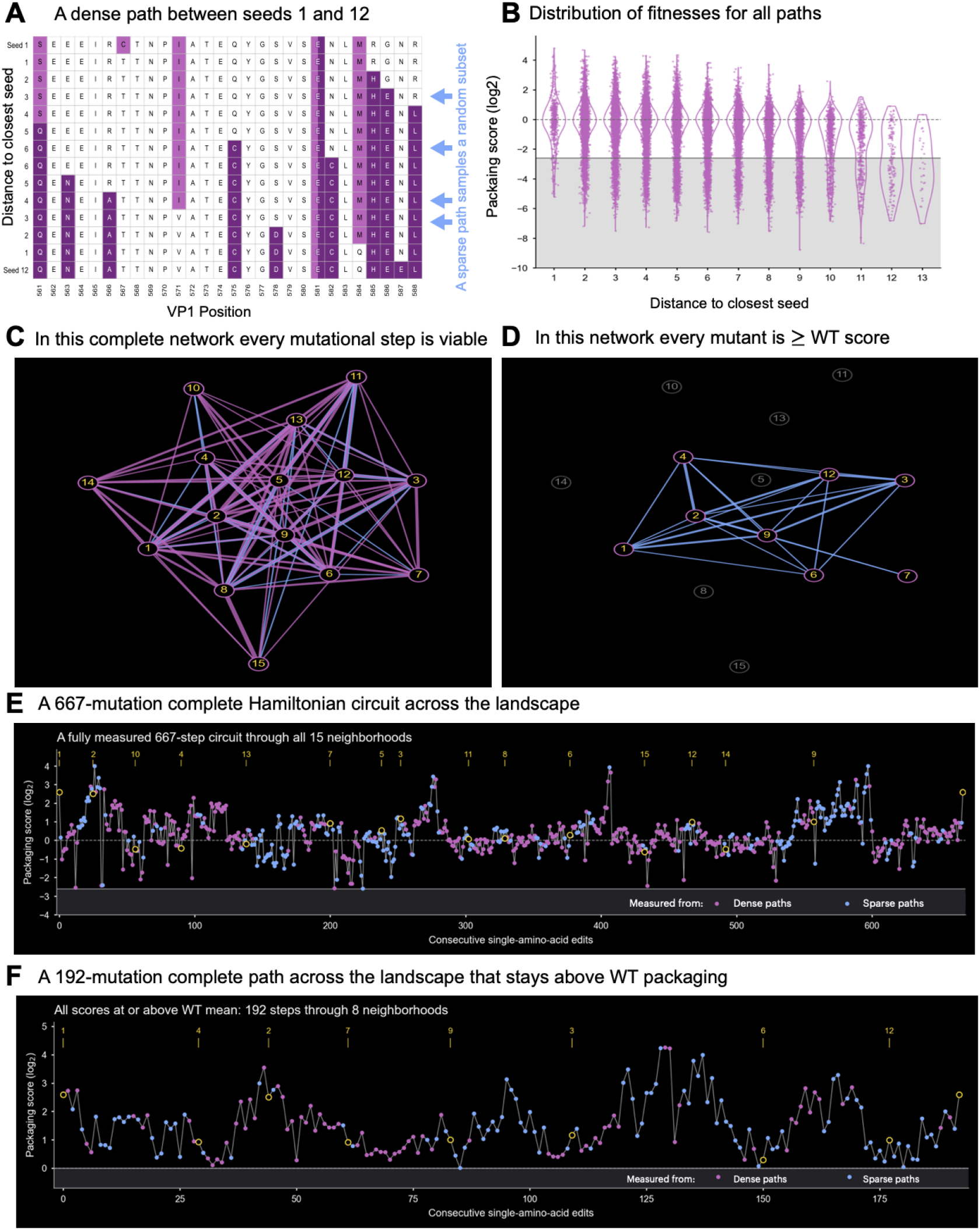
Billions of long function-preserving Hamiltonian cycles are possible in this measured fitness landscape. A: An example of a dense path between two seeds, a sparse path would skip some steps. Colored cells show mutations to the WT reference. B: Distribution of fitnesses of all complete and sparse paths plotted against the sample distance to the closest seed. C: A network of completely viable neutral paths among the seeds (where every single-step mutation is viable), with wider sets of edges between two seeds, meaning more distinct paths were found. Paths consisting entirely of dense samples are shown in pink, paths that were constructed by combining dense and sparse samples are shown in blue D: A subnetwork where all the mutants are at or above WT in packaging competence. E: An example of a viable Hamiltonian circuit of a total length of 667 mutations we found in this dataset, visiting every seed once and traversing every one. F: A circuit in the subnetwork with neutral or better than WT packaging.

We find that pairwise stochastic paths, without the use of ML models, yield strikingly robust viability, recovering functional ridges crossing the landscape (Fig. 4B, C). Of 315 densely sampled paths, 183 were complete and fully evaluable after experimental quality control; 85 of these retained viability at every measured step. Combining dense and sparse path samples reveals a large viable network connecting all 15 seed neighborhoods (Fig. 4C). Within the network built entirely from measured variants, we enumerate upward of 10⁹ distinct Hamiltonian circuits (5.4 × 10⁹ for the measured viability calls, and no fewer than 2 × 10⁹ in any bootstrap over sequencing replicates or resampling of read counts, see Fig. S6), each visiting every seed once before returning to its start. These circuits span 212-667 single amino-acid steps within the network, with every intermediate retaining packaging competence, and represent roughly 6% of all possible orderings within the 15 neighborhoods. Figure 4E shows a 667-step example. Connectivity remains under a much stricter requirement: eight neighborhoods remain connected where every measurement is at or above mean WT packaging (Fig. 4D). These circuits span 116-192 single amino-acid steps, including the 192-step example shown in Figure 4F. Thus, extensive navigation is possible not only while retaining viability, but also while maintaining measured packaging at or above the WT reference. This result reveals substantial latent capacity for functional diversification within this region of the capsid. Extensive neutral networks have long been predicted theoretically and observed experimentally but have not been demonstrated in proteins across comparably vast volume of the sequence space (Rotrattanadumrong and Yokobayashi 2022; Klug et al. 2019; Greenbury et al. 2022; Schuster and Stadler 2023; Poelwijk et al. 2019). This connectivity provides potential routes for extensive sequence exploration through both selection and genetic drift, provided fitness differences along these paths are sufficiently small. In this regime, network topology can become an important determinant of evolutionary outcomes, influencing which genotypes are favored and how mutational robustness evolves (Wagner 2008; van Nimwegen et al. 1999).

### Models discover regions that are hard to reach for evolution

We use machine learning guided walks using the same classification models in (Bryant et al. 2021). We use the exact same models, training data and the same weights: Logistic Regression (LR), Convolutional Neural Networks (CNN), and Recurrent Neural Networks (RNN) models were trained on three distinct datasets around WT with (i) all singles and random double mutations (CR2, n=2,868) (ii) all singles and random multi-mutations up to 10 edits to WT (CR10, n=9,020) (iii) Random mutants up to distance 10 and additively designed mutations up to edit distance 39 (A39, n=64,280), from which the 15 neighborhoods were selected (see methods here and in (Bryant et al. 2021) for details). These models are not trained on evolutionary priors, only assay data, which enable them to steer independently from the evolutionary record of AAV2.

The main difference between model-guided design and the other ways we sampled this landscape is the type of mutations neural models introduce (Fig 5A). Random local mutants are 42.8% viable one edit from a seed and only 1.6% viable at five, and past that distance we measure essentially nothing that packages. Stochastic paths stay viable almost by construction, each step is selected to be viable in at least one background, and the recombinant paths we could follow run out at around 13 edits to the closest seed. While model-guided designs start below the stochastic paths in terms of overall viability, they attempt more ambitious (non-viable) changes and decay slowly. While these models succeed at predicting viability at far distances they do not generate capsids with peak performance: the highest measured fitnesses were sampled as walks or random mutations one or two mutations from a seed, and 46 of the top 50 best producers were derived from seed 2, the most epistatic seed.

**Figure 5:**
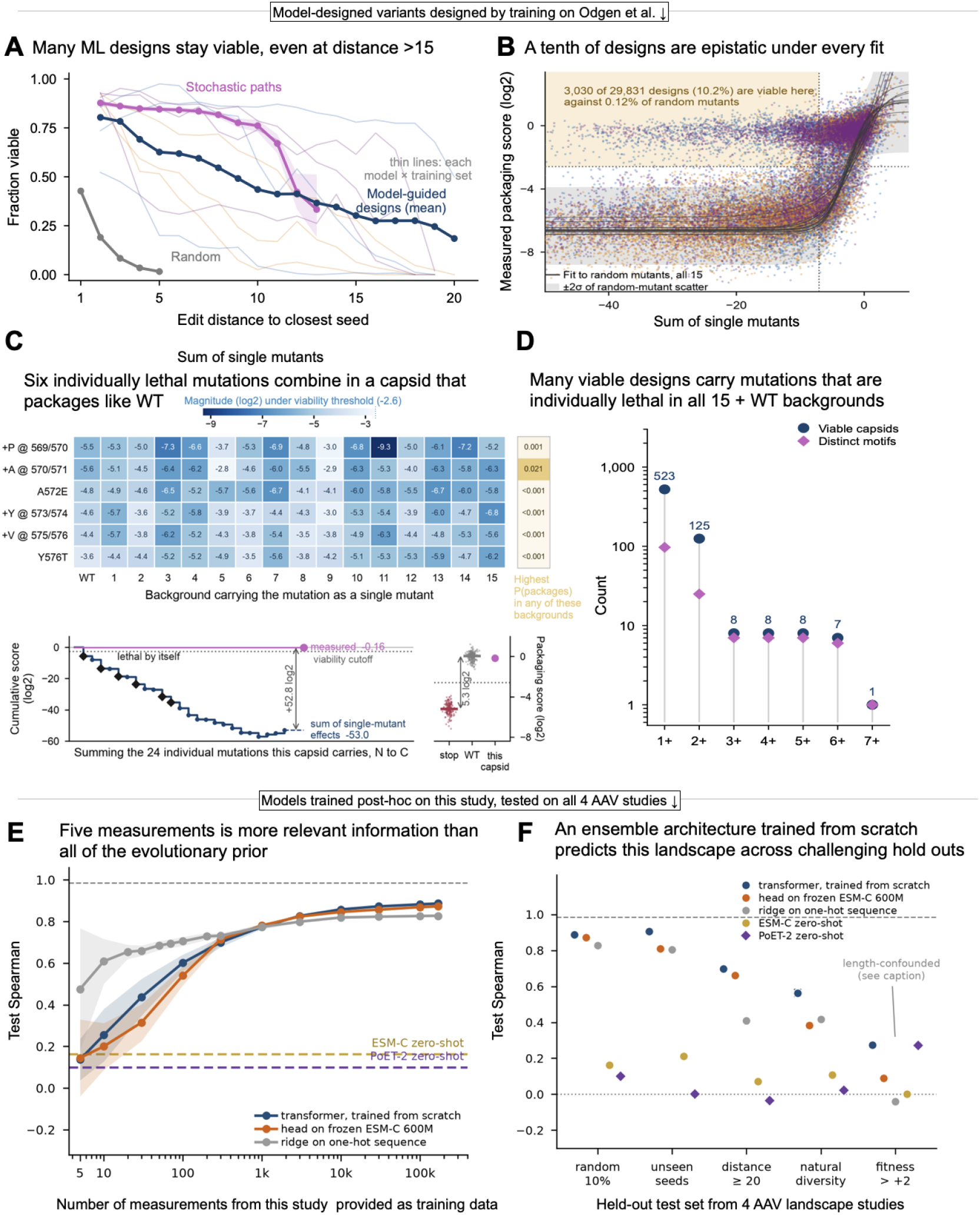
Model-guided design reaches deep into the sequence space and finds combinations that are difficult for evolution and recombination to find. A: Fraction viable against edit distance to the nearest seed, for random local mutants, neutral paths and model-guided designs B: Measured packaging score against the sum of single-mutant effects for 29,831 designs in the fifteen neighborhoods. Thin lines are the fifteen global epistasis fits (per neighborhood), each fitted on held-out random mutants separately C: Top, the six universal lethals carried by one viable capsid (24 edits from wild type), measured as single mutants in wild type and in each seed; all 96 values fall between −9.30 and −2.81 log2 (white marks the cutoff). The right column gives each mutation’s probability of packaging in its most favourable background, at most 0.021, and ∼10⁻³⁴ for the six together, from a logistic calibrated on the assay’s stop and wild-type controls. Bottom, cumulative sum of the capsid’s 24 measured single-mutant effects, N to C. D: Viable capsids introducing k or more universal lethals: single mutants below the cutoff as in wild type and in every seed where they were measured counted as distinct capsids and as distinct motifs. {E,F} We use this experiment’s data to predict held out sequences from all 4 AAV libraries measured in different studies. E: Test spearman results for from-scratch training of a baseline ridge regression and an ensemble of transformers compared to zero shot and partially trained predictions from models trained on evolutionary data. F: Comparison of the performance of these models on 5 qualitatively different heldout tasks. Predictions on fitness > +2 category are in part explained by the number of insertions (see Table S3).

The training data matters more than the model architecture, with Logistic Regression being the most sensitive to limited input data. Comparing designs at five edits from their designated parent, every (architecture, training set) pairing is enriched over random mutants, by 10- to 61- fold in viability hit rate. Models trained on CR2 are not uniformly bad at predicting viability; they rank adversarial examples, sequences that score well at design time and measure poorly highly, which produces a designed batch full of false positives. This is the behaviour expected of model-guided design when the design distribution sits far outside the training distribution, termed distribution shift (Quinonero-Candela et al. 2022). As in (Bryant et al. 2021), LR trained on the standard dataset (CR10) is the best performer at introducing mutations without breaking the capsid.

Aggregate statistics still paint an incomplete picture; they do not reveal whether a design is reachable by adding up what we already know about its parts. Fitting the global epistasis model of each neighborhood to random mutants and applying it to the designs of that neighborhood, the designs depart from the global epistasis model in one direction: they are viable where the fit predicts they wouldn’t be. The global epistasis fits across backgrounds are close enough to one another that we hold all designs to the same conservative standard; the envelope of all fifteen curves widened by twice the scatter of random mutants about their own fit. Nonetheless, we find 3,030 of 29,831 designs (10.2 %) measured viable where that widened family predicts they wouldn’t be, against 0.12% of held out random mutants judged the same way (Fig 5B). If we instead judge each design against its own neighborhood’s fit at the viability cutoff, without that margin, the departure rate is 22.6% for designs and 2.1 % for random mutants, and designs exceed the random rate in all fifteen neighborhoods (Fig 5D). The effect is not explained by distance: designs restricted to two to five edits from their parent are viable at an 11.0 % rate. RNN-optimized samples most often generate variants that should not have been packaged according to the additive model.

As an example, one RNN-CR10 designed viable capsid (Fig 5C), at 24 edits from wild type, carries six mutations that are individually *universally* lethal. Each individual mutation was sufficient to break packaging in wild type and in every one of the fifteen seed backgrounds (Fig S4), yet the design contains multiple of them. Calibrating packaging score against the controls at its most favourable measurement in any background (from all 15 + WT), the best of the six mutations has a 2.1% chance of packaging and four of them are below 0.1%. An independent-hits model for these mutants would assign the probability of a viable combination at ∼10^-34^. Notably, none of these motifs appear in the multiple sequence alignment of dependoparvoviruses (310 aligned sequences), no insertions are observed in this segment naturally, and substitutions at positions 572, and 576 do not contain E or T respectively, and rarely contain amino-acids with similar chemical properties.

We find many variants like these (Fig 5D). Across the library, 523 viable capsids introduce at least one universal lethal mutation, 125 introduce at least two, 8 introduce at least three, and one carries seven. Because designs from the same walk share mutations, we also count distinct sets of universal lethals. About half of these variants come from RNN-based designs, which in general have 2.2-3.4 times higher per-design success for capsids containing lethal mutations than the next best architecture (0.38%, CNN).

### A vast space of functional variants can be discovered by a small amount of local data, but not by general-purpose models

Finally, we study the broad structure of this landscape by combining the measurements of all four major AAV studies in this sequence region (Ogden et al. 2019; Bryant et al. 2021; Sinai et al. 2021), which leaves us with 65945 + 208023 + 212558 +7290 = 493816 unique measured proteins from this space. We benchmakr two leading protein sequence representation models trained on evolutionary background, ESM-C 600M (Candido et al. 2026) and PoET-2 (Truong and Bepler 2025). In addition, we train a simple ridge regression, and an ensemble of small transformers from scratch on one-hot sequences, as well as training a regression head on top of the ESM-C representation. We find that with as few as five samples from this landscape, ridge regression can predict random held out variants better than more complex model, fine-tuned or zero shot. With more data added, complex architectures overtake ridge as a predictor, but the evolutionary prior does not add a distinct benefit. Around one thousand examples (0.6% of the data) is sufficient to learn close to 90% of ceiling performance achieved by each architecture. We repeat the test by constructing 5 distributionally distinct target sets for prediction from all 4 experiments, trained on subsets of sequences in this study. The target sets include a random 10% hold out set, 3 held out seeds and their neighboring variants, held-out variants at edit-distance above 20, variants generated by importance sampling the AAV MSA background, and variants with packaging score above 2 (see methods section 4.2). This pattern persists across test sets. PoET-2 is an exception on the high-fitness holdout data, although its apparent advantage disappears after controlling for insertion count.

The combinatorial residue space attempted in this study contains ∼10^36^ sequences if we only consider substitutions, and ∼10^75^ if we account for insertions as well. Excluding mutations in the universal lethal set leaves ∼10^33^ substitution sequences as possibly viable, and a more conservative estimate accounting for the fraction of viable variants for each substitution still leaves ∼10^27^ variants as possibly dense in viable proteins under these simple assumptions. We do not claim that the measurements in this study generalize readily to this entire space, but as shown in Fig 2A, the span of sequences we cover is large, and every model that learned to avoid the universal lethals generated variants up to 10 edits from a starting point (which is not particularly unique as a sequence) with viability rate near 50%. The universal-lethal set defines the hull of a subspace containing landscape orebodies. Leave-one-seed-out analysis shows that these simple rules generalize: without the held-out seed, presence of these mutations predicts the viability of random variants around the left out seed with AUROC 0.868 (and almost no loss in performance compared to the full set). Notably, it is asymmetric, i.e. carrying a universal lethal mutation is a good exclusion rule (NPV 0.995, sensitivity 0.985), whereas passing the filter raises the density of viables within a landscape subspace (PPV 0.489). The highly viable graph of the previous section is a narrow ridge exploring this orebody. Across the four chips, 51.3% of the 493,816 measured variants are viable. Of all viable variants, 98.0% fall within the hull, including every one of the 1,202 viable variants among the 6,184 sampled randomly at twenty or more mutations from wild type. The majority of our results are qualitatively consistent with recent theory predicting that the upper reaches of additive (and in some cases global epistatic) fitness landscapes are blunt rather than sharply peaked, with a correspondingly large supply of high-fitness genotypes (Kinney 2026), although the neighborhood around seed 2 is relatively sharp (and comparatively non-additive). We view the direction of surprise in this study to be one that denotes the landscape as more permissive to mutations than it would appear from a single deep mutational scan around the wildtype.

## Discussion

We make several new observations through an expansive, quantitative sampling of random and designed variants within the sequence space of AAV2 capsid protein. First, additive and global epistasis patterns are universal in this sequence space, but the degree to which these patterns explain local structure is different among neighborhoods. Much of the apparent epistasis may be a function of the fitness-epistasis uncertainty principle (D. Brookes et al. 2024): concentration of the observed fitness function implies a dense epistatic representation, even when the underlying latent function is sparse. Consistent with this picture, accounting for global epistasis leaves little additional contribution from explicit pairwise interactions. Beyond this correction, we highlight both idiosyncratic mutations, where they act highly differently depending on the background they are in and idiosyncratic neighborhoods, where self-prediction is far better than inbound and outbound prediction. We also show that viability decays quickly, and is largely governed by the arrival of single mutations that are lethal, rather than accumulation of mildly deleterious mutations.

Second, we observe that stochastic recombinant paths are extremely robust and use them to find a network of densely connected neutral paths between our original seeds, suggesting AAV capsids may have many paths available to evolve away from any antibodies that target this region or to discover new functions. This is surprising in the context where about half of single mutations (in some cases up to 80%) would break the capsid and almost all random mutants at distances above 3 to any seed are non-viable. The overall fitness of the capsid is composed of multiple relevant properties and many of these mutations are likely pleiotropic, which would reduce the space of naturally or therapeutically relevant viruses significantly (Weinreich et al. 2006). However, while the structure of fitness landscapes remains highly heterogeneous (Bank et al. 2016) our results match and expand the emerging picture of empirical fitness landscapes from other proteins. We see that the protein landscape can be rugged yet navigable at short distances (Papkou et al. 2023), and many functional proteins may exist in the unobserved sequence space (Diplock et al. 2026; Avizemer et al. 2025), even if many direct paths to them have not been explored. This study sits between these regimes: within a highly epistatic and overwhelmingly non-viable landscape, we empirically find complete and long mutational paths that form a large connected network spanning distant viable capsids that offer the protein many paths to evolve, but also contain many viable paths that are not easily evolvable.

Third, we show that deep exploration of the landscape, away from the additive manifold and the evolutionary record, is possible both through independent sites and more complex models. Independent-site models may get there by first changing the background enough with safe mutations, and then adding new mutations that, in the new context, would be better tolerated. This is a route also accessible to evolutionary search, through mutations that side-step fitness valleys (Oros and Krug 2026) or increase evolvability. In contrast, more complex models like the RNN offer a unique (and likely distinct from evolutionary search, given the relative evolutionary inaccessibility) tool for protein design. Thinking about the performance of these methods for sampling a fitness landscape depends on the objective. AAVs are traditionally selected for a particular function (Huang et al. 2024; Tabebordbar et al. 2021), which is an effective development approach but leaves much less control over how the objective is achieved. Our results suggest that the landscape structure itself offers hints of how to sample it while maintaining control and efficiency. There is plenty of additive structure but epistasis can also be exploited to achieve performance: designing very high-performing proteins is easier to achieve if there is a high-performing and rugged seed neighborhood to sample from (e.g. Seed 2 in our case), and even 1-2 mutations may be sufficient to improve the protein (Jiang et al. 2024). Designing sequences at high edit distance (Freschlin et al. 2024; Bryant et al. 2021), or those that are hard to discover by evolution, requires using models that have learned some implicit information about the landscape’s behavior overall, for instance, to learn and avoid lethal mutations.

Fourth, a global boundary, largely inferrable from a few lethal variants, contains the majority of the functional observations in this study. Avoiding universally lethal variants from local measurements leaves a massive space open to exploration. Moving within this space changes the shape of the boundary, but is hard to achieve through additive models learned in some local subspace. We see a few successful examples of local non-additive models managing to step outside this boundary, as discussed in the previous paragraph. More broadly, if this structure (single-hit + additive) is common in the protein space, it could point to a simple generalization rule, likely learnable by foundation models: much functionality can be simply predicted by learning what mutations to avoid, introducing a few clearly tolerated mutations, and gambling on mutations that aren’t clearly in the banned list. More broadly, while the sequence space is huge and functional proteins may be a very small fraction of it, there are astronomically many functional proteins that can be found using simple rules learned from local structure. The apparent success of sequence-based machine learning models trained on evolutionary data may reflect the landscape’s permissiveness more than the power of the ML architecture or sophisticated learning as the necessary information is simply not in the evolutionary data. Beyond sequence-only models, currently structure prediction models show comparatively low ability to predict variation at the single-mutation level far from WT, in particular for unstructured loops like those studied here. Therefore additional accuracy in modeling of a local landscape by general models is unlikely without supervised fine-tuning with local data, as shown by the private information retained within each background.

Today, a majority of biological sequence models are trained mostly on evolutionary data. Not only is the collective human sampling of evolutionary data highly biased (F. Ding and Steinhardt 2024), the evolutionary record itself is a highly biased and narrow sample of the functional space. While evolutionary information can and does enable the discovery of many valuable protein sequences (Russ et al. 2020) sequences explored by evolution are the result of sampling through selective and stochastic forces that are suboptimal from the perspective of learning a landscape structure and have no inherent alignment with human interest. In fact, one would expect that proteins which survive evolutionary history would not be those that have highly fragile, unnavigable landscapes, as it would make them difficult to evolve. Nonetheless, they may be highly valuable for human use. What we found by models built on deep mutational scans is akin to prospecting signals about the regions of sequence space, revealing landscape orebodies, that likely include a high number of functional and useful proteins, which would not be found from in silico evolutionary priors or even empirical directed evolution campaigns alone. This study shows that collecting even a small amount of data can teach us highly usable, simple insights on the local structure of the landscape which can then be used for design. We anticipate that exploring the space of functional variants for this (and other) protein spaces beyond evolution’s reach should provide vast reservoirs of molecules with high therapeutic potential.

## Materials and Methods

### 1. AAV mutant sequence library generation and production assay

We constructed libraries using a method similar to that previously described by Ogden et al (2019) and Bryant et al. (2021). The following protocol is identical to that of Bryant et al. (2021) and the following is replicated from that paper (experiments were done concurrently using the same methods in separate pools). For validation experiments, 184-mer DNA oligonucleotides (oligos) were synthesized as single-stranded DNA by Agilent. Designed amino acid sequences were back-translated to nucleotide sequences by choosing any possible codon (generally keeping the WT codon and selecting mutant amino acid codons with no bias, but disallowing codon choices that created restriction enzyme sites used in cloning). From 5′ to 3′, each oligo contained a forward primer binding site, a BbsI restriction site (5′-GAAGACAT|TACA-3′), a nucleotide mutant coding sequence (with > 83 nucleotides), a BsaI restriction site (5′-CAAG|CGAGACC-3′), an EcoRV ‘kill-cutter’ restriction site, a BsaI restriction site in the opposite orientation (5′-GGTCTCA|CGCT-3′), an 18-nucleotide barcode sequence, a BbsI site in the opposite orientation (5′-CGCT|AAGTCTTC-3′) and a reverse primer binding site. Barcodes were included in the synthesized oligos but were not used for downstream sequencing or analysis (rather, the mutant coding region was directly amplified and sequenced, matching the amplification method used in the previous production experiments). PyDNA32 was used for in silico testing of the cloning process, ensuring sequence compatibility with the cloning strategy. The code for executing this process is provided in the synthesis pipeline component of the bioinformatics pipeline code. An example oligo with WT coding region is 5′-GGGTCACGCGTAGGAGAAGACATTACAGACGAAGAGGAAATCAGGACAACCAATCCCGTGGCTACGGAGCAGTATGGTTCTGTATCTACCAACCTCCAGAGAGGCAACAGACAAGCGAGACCGATATCGGTCTCACGCTGTAATGCGGTCTGAGCCGCGCTAAGTCTTCGTGTGGCTGCGGAAC-3′.

Cloning was carried out in three steps. First, oligos were PCR amplified using Q5 high-fidelity DNA polymerase (NEB, no. M0492) and an annealing temperature of 60 °C (forward: 5′-GGGTCACGCGTAGGA-3′, reverse: 5′-GTTCCGCAGCCACAC-3′). A backbone plasmid containing the WT AAV2 cap gene was also amplified with Q5 and an annealing temperature of 72 °C (forward: 5′-TTGGTCTCA|CGCTAGAGACGGTGTGGCTGCGGAAC-3′, reverse: 5′-AAGGTCTCC|TGTAATCATGACCTTTTCAATGTCCACATTTG-3′). This PCR was used to add

BsaI sites with overhangs complementary to BbsI overhangs in the oligos (as indicated by cut sites in primer sequences). Amplified oligos were digested with BbsI-HF (NEB, no. R3539), and amplified plasmid was digested with BsaIHF-v2 (NEB, no. R3733) in separate 50-µl reactions. Digest products were purified using homemade SPRI beads, mixed at a 3/1 molar ratio (oligos/plasmid) and ligated using T4 DNA ligase (2×106Uml–1; NEB, no. M0202). Ligation products were ethanol precipitated and transformed into 50µl of electrocompetent cells (Lucigen 10G SUPREME, no. 60081). Following 1-h recovery at 37 °C, cells were added to 4ml of selection medium (2× YT with kanamycin) and grown at 37 °C overnight. The following morning, step1 library plasmids were mini-prepped by alkaline lysis (Qiagen, no. 27104). In this first cloning step, oligo sequences replaced the corresponding 84-base-pair WT sequence and the 3′ region of the cap gene in the backbone plasmid. In the second cloning step, an amplicon containing the 3′WT region of the cap gene was generated from the initial backbone plasmid using Q5 and an annealing temperature of 72 °C (forward: 5′-TTGGTCTC A|CAAGCAGCTACCGCAGATGTCA-3′,reverse:5′-AAGGTCTCA|AGCGAGAGACGTCCTACGCGT GACCC-3′). This amplification step was also used to add BsaI sites complementary to those in the oligos. Step1 library plasmids and the 3′WT amplicon were separately digested with BsaI-HFv2, bead purified and ligated as above. Ligation products were digested with EcoRV-HF (NEB, no. R3195) in a kill-cutting step to remove step1 plasmids that did not incorporate the 3′WT amplicon. EcoRV digest products were ethanol precipitated, transformed and mini-prepped as in step1. In the third cloning step, a destination ITR-containing plasmid was digested with HindIII-HF (NEB, no. R3104) and SpeI-HF (NEB, no. R3133). Complete mutant cap gene sequences were amplified from the step2 plasmid library using Q5 and an annealing temperature of 70 °C (forward: 5′-AGGTCTCA|AGCTTCGATCAACTACGCAGACAG-3′, reverse: 5′-AGGTCTCA|CTAGATGAGCTCGTCGACGTTCC-3′). This amplification step was also used to add BsaI sites and overhangs complementary to the HindIII and SpeI sites in the ITR plasmid. Amplicons were digested with BsaI-HFv2 as above. Digested ITR plasmid and step2 library amplicons were bead purified, ligated, transformed and mini-prepped to generate the final plasmid library. For the earlier rounds of library cloning, creation of mutant cap genes was accomplished in a single cloning step since oligos did not contain BsaI sites, EcoRV sites or barcode sequences, enabling cloning directly into the corresponding position in the WT cap gene. Ligation sites and oligo and cap PCR primers for this single-step cloning were the same as above. Similarly, the final library cloning step to move the mutant cap gene sequences into the ITR plasmid remained the same. The final plasmid library was transfected into HEK293T cells to produce viral particles. Cells were grown in DMEM (ThermoFisher, no. 10566016) supplemented with 10% fetal bovine serum (ThermoFisher, no. 10082147) and seeded in five-layer cell stacks (Corning, no. 353144) 2d before transfection. Polyethylenimine (PEI) was used for transfection at a mass ratio of 3/1; 125ug of adenovirus pHelper plasmid, 75ug of an AAV rep plasmid and 1ug of library plasmids were mixed with PEI, incubated for 20min and added to cells. Media were changed completely at the time of transfection and replicate transfections were carried out in separate cell stacks. Here, the lower levels of library plasmid were chosen to reduce the number of plasmids transfected into individual cells, such that the potential for mosaic capsid formation and cross-packaging was minimized. Three days post-transfection, 5M NaCl was added to the cultures for a final concentration of 0.5M and cultures were incubated at 37 °C for 3h. Following incubation, mixtures were transferred to fresh containers and incubated at 4 °C overnight. The next day, the resulting supernatants were run through 0.22-µm PES filters (Corning, no. 431098); 40% PEG-8000 was then added to give a final concentration of 8%, and mixtures were incubated at 4 °C for 3h. Samples were centrifuged at 3,000g for 20min to pellet the PEG precipitate, and pellets were resuspended in 7ml of DPBS. Viral genomes external to the capsid and carryover plasmid DNA were degraded with benzonase; a 10,000-fold dilution in benzonase (Millipore Sigma, no. 1.01695.0001) was added to resuspended pellets, and samples were incubated at 37 °C for 45min. Encapsidated genomes were separated from the remaining cellular debris using iodixanol ultracentrifugation and concentration via size-exclusion spin filters as described previously 19,33. Briefly, benzonase-treated samples were underlaid with an iodixanol gradient (Sigma, no. D1556) in polypropylene tubes (Beckman Coulter, no. 362183) and centrifuged at 242,000g for 1h at 16 °C. Capsids were collected from the 40% iodixanol fraction and concentrated using a spin concentrator (Millipore Sigma, no. UFC910024) to generate the final purified pool. Cap gene sequences remaining in the purified pool represent mutants viable for capsid assembly and genome packaging. Purified capsids were heat denatured at 98 °C for 10min, and PCR was run with Q5 and an annealing temperature of 65 °C to amplify the mutant region of the cap gene (forward:5′-GCTCAGAGAAAACAAATGTGGAC-3′,reverse:5′-GAACGCCTTGTGTGTTGACATC-3′). PCR reactions were carried out in the presence of EvaGreen (Biotium, no. 31000) and run on a BioRad CFX96 quantitative PCR machine to ensure that reactions were stopped during the exponential phase. Illumina sequencing adapters and indices were added in a subsequent PCR. These PCR amplicons were sequenced with overlapping paired-end reads using Illumina NextSeq. Paired-end reads were merged to generate a consensus read using PEAR34, and read counts were calculated for every member of the designed library. Reads with a minimum Qscore of 20 were selected for four technical plasmid replicates and three biological virus replicates (each with at least two technical replicates; Supplementary Fig. 2). Mutant fitness in the viral production assay was calculated by taking the ratio of mutant read counts in the viral library over the counts in the original DNA library, normalizing by the ratio of the WT sequence. Measurement of viral genome abundances from tissues for the design of the A39 dataset was done via amplicon sequencing from purified vector genomes, with PCR protocols as described above. Three separate batches of virus were prepared and 3.5×10^10^ (batch 1), 2.3×10^10^ (batch 2) and 3.5×10^10^ viral genomes (batch 3) of the C1 virus library were diluted in 200 µl of PBS and injected into mice, four mice per batch, for 12 mice in total. Mice were all 8 weeks old, male and C57BL/6J. For each of the three batches, two mice were injected retro-orbitally and two intraperitoneally. After 1, 5 and 24h, 30 µl of blood was drawn by facial bleed and frozen on dry ice, then at −80 °C. After 8d, mice were dissected and tissue samples from liver, kidney, heart, lung, brain, spleen, muscle, skin, stomach and testis were frozen on dry ice, then at −80 °C. Approximately 150mg of each organ was ground using a disposable mortar and pestle (Kimble Chase, no. 749625-0010). DNA was purified from tissues using alkaline lysis (Qiagen, no. 27104), and from blood using the Qiagen MinElute Virus Spin Kit (no. 57704). Biodistribution was similar across both routes of administration. The overall effect on biodistribution of viral genomes for each organ and blood sample was calculated in R using DESeq2 across measurements from multiple mice, combining 12mice for blood and liver and four mice from batch2 for the remaining organs. The animal protocol was approved by the Harvard Medical School Institutional Animal Care and Use Committee. The A39 Library was used to select seeds for this study. Seeds were selected based on their packaging viability, as well as liver biodistribution in mice.

## 2. Design methods

### 2.1 Sequence representation

Sequences are one-hot encoded on a 21-state × 56-coordinate grid (21 states = 20 amino acids plus a stop symbol (denoted by *); 56 coordinates = two per residue of the 28-residue region, so that single-residue insertions can be represented). Flattening this encoding gives the 1,176-dimensional input vector.

### 2.2 Construction of ML training datasets CR2, CR10 and A39

The architecture of the neural network models and the training data discussed in this section is identical to (Bryant et al. 2021) and is included for easy referencing.

The experimental design involved constructing three distinct training datasets, each containing sequence variants sampled from a constrained sequence space around the wild-type AAV2 sequence. These datasets were generated using different sampling strategies to explore various regions of the fitness landscape and evaluate model performance effectively.

1. **CR2 Dataset (N=2,868, 40% Viable)**: This dataset combines:

a. **C:** The complete set of single-site mutants, comprising 1,112 variants with 58% viability. Single mutants include all possible substitutions across 28 residue positions, as well as all possible single-residue insertions at 29 insertion positions. This results in 1,112 single-site mutants.
b. **R2**: A random subset (<1%) of the possible double mutants, comprising 1,756 variants with 29% viability.
2. **CR10 Dataset (N=9,020, 16% Viable)**: This dataset includes:

a. **C**: The same 1,112 single-site mutants described above (58% viable).
b. **R10**: A set of 7,908 randomly generated mutants containing 2–10 mutations, with 10% viability. These randomly generated mutants are fully distinct from the validation set used for model evaluation. Despite the low viability of these variants, they provide valuable information for training ML models to distinguish functional from non-functional sequences.
3. **A39 Dataset (N=64,280, 56% Viable)**: This dataset consists of:

a. **A39**: A baseline additive dataset containing 56,372 variants generated using a single-site additive fitness model, with a viability of 62%.
b. **R10**: The same 7,908 randomly generated multi-mutants described in C1+R10 (10% viable).

Importantly, R10+A39 does not include the 1,112 single-site mutants from C1. The inclusion of A39 allows for a direct comparison of datasets that either explicitly include all single mutants (C1+R10) or emphasize higher-order variants derived from additive models (R10+A39).

For all three datasets, only variants with a plasmid count >100 were included to minimize noise in fitness measurements caused by low plasmid counts. Each dataset includes at least four synonymous nucleotide sequences for every amino acid sequence variant, and the highest observed fitness measurement across synonymous variants was used to represent each unique amino acid sequence.

### 2.3 ML Model Experimental Design Overview

All three datasets were used to train classification models that predict whether a given AAV2 capsid variant is functional (Fig. 1b). Three types of model architectures were employed:

**1. Logistic Regression (LR):**

A baseline model that does not capture higher-order mutational interactions. The simplicity of LR models reduces the risk of overfitting, particularly when trained on smaller datasets, at the expense of predictive complexity.

**2. Convolutional Neural Networks (CNNs):**

These models capture contiguous local mutational patterns as raw features and aggregate them into larger receptive fields at deeper layers.

**3. Recurrent Neural Networks (RNNs):**

Specifically, unidirectional, multi-layered long short-term memory (LSTM) networks were used to aggregate knowledge of mutations along the sequence in a stateful manner.

All models used a one-hot encoding for input sequence representation, with supervised learning based on binary labels indicating capsid functionality.

### 2.4 Training ML Models For design

The cross-product of three model architectures (LR, CNN, RNN) and three datasets (CR2, CR10, A39) yielded nine combinations of models (e.g., LR{CR2}, RNN{A39}). Each combination involved training 11 replicas with different random initializations to create ensemble models. Model performance, evaluated on a held-out validation set of randomly sampled mutants, was used for early stopping. The ensemble’s mean replica score was used to rank sequences for model-based selection and design.

**Optimization**: CNN and RNN models were trained using the Adam optimizer, with learning rates of 0.001 and 0.01, respectively. LR models were optimized using the FTRL algorithm with a learning rate of 0.01. Learning rates were selected via a hyperparameter sweep based on validation performance using the C1+R10 dataset.

**Regularization**: Early stopping which halted training after 10 evaluation periods without improvement in validation precision. Each evaluation period comprised 500 steps, with a batch size of 25. Training wall-times ranged from 20.3 to 85.3 minutes

### 2.5 Architecture Selection and Hyperparameter Tuning

Neural network architectures varied in complexity:

- **Logistic Regression (LR)**: A simple model with 1,161 parameters and no hidden layers.
- **Convolutional Neural Network (CNN)**: A model with 55,189 parameters and four hidden layers: Input shape: (58, 21), Two convolutional layers (7-width, depths of 12 and 24) with pooling and batch normalization. Two fully connected (FC) layers with batch normalization.
- **Recurrent Neural Network (RNN)**: A multi-layer LSTM model with 128,901 parameters and two hidden layers, each containing 100 units. Input shape: (21 tokens). Two stacked LSTM layers.

All hyperparameter tuning was conducted using the C1+R10 dataset and evaluated on a disjoint validation set (N=1,977). Binary softmax cross-entropy loss was used for all models.

## 3. Descriptive Local Models

### 3.1 Additive and global epistasis models

We build descriptive models of the quantitative behaviour of the landscape around each seed. The absence of an insertion is encoded explicitly in the otherwise-unused stop channel at insertion coordinates.

### 3.2 Local (additive) model

For each seed *i* we assemble the measured single insertion and substitution effects into a matrix *Mᵢ*, each effect obtained by subtracting the independently calibrated prior-assay seed score. Given a variant *x* with one-hot encoding hₓ, its additive fitness is *fₐ(x; i)* = *cᵢ + Σ (Mᵢ* ∘ *mᵢ* ∘ *hₓ)*, where ∘ is the Hadamard product, *Σ* the sum over matrix elements, *cᵢ* the seed baseline, and *mᵢ* a mask that is zero at the residues carried by seed *i* and one elsewhere, so only the mutations *x* carries relative to the seed contribute.

Alternatives with no measured single-mutant effect are imputed by the median measured effect at that coordinate within the training seed, falling back to the overall training-seed median. This model has no free parameters; we refer to it as the local model. We additionally construct a WT-centric additive model, fₐ,WT, which scores the same seed-relative mutations using the matrix of single-mutant effects measured around the wild type.

### 3.3 Global-epistasis model

We elaborate the additive model with a monotonic global non-linearity applied to the additive trait, gᵢ(x) = σ(fₐ(x; i)), where σ is a 4-parameter increasing logistic *σ(z) = L / (1+ exp{-k(z − z_0_) })+ b*. For each seed the parameters are fit by non-linear least squares (scipy.optimize.curve_fit) using the non held out multi-mutants in that seed’s training set (these were not refit against a test score).

### 3.4 Cross-prediction benchmark

A source model was trained on each of the 15 seed neighbourhoods. Local performance used the model trained in that neighbourhood; transfer performance was the mean over the 14 other source neighbourhoods evaluated on the same target test set. We report Pearson *r*, *R²*, and RMSE.

### 3.5 Ridge regression

Ridge regression (scikit-learn, https://scikit-learn.org) used the same one-hot encoding, centred on the training seed, plus an intercept, with the target standardized by the training mean and standard deviation. The L2 regularization coefficient was selected from {0.01, 0.1, 1, 10, 100} by validation mean-squared error.

## 4. Full Landscape Models

We combine sequence data from 4 publications: Ogden et al 2019. (additive and random designs), Bryant et al 2021. (LR/CNN/RNN walks and seeds), Sinai et al 2021. (VAE and importance-sampled alignment designs), and this study (LR/CNN/RNN/sparse/dense paths, random viable variants, and per-seed variants)

### 4.1 Models

#### 4.1.1 From-scratch transformer training

Insertion-aware tokenisation: 20 amino-acid tokens plus padding, a separate insertion-indicator embedding so an inserted residue is distinguishable from a substituted one at the same index, and a learned positional embedding; MAXLEN 43.

Encoder: 6 × `TransformerEncoderLayer`, d_model 192, 6 heads, feed-forward 384, dropout 0.1, pre-layer-norm, GELU. Masked mean-pooling over non-pad positions → LayerNorm → head `Linear(192,192) → GELU → Linear(192,1)`, output clamped to ±20. 1,832,457 parameters.

Training: AdamW, lr 3 × 10⁻⁴, weight decay 0.01, batch 512, 15,000 steps, 750 warm-up steps then cosine decay, gradient clipping 1.0. Validation Spearman evaluated every 750 steps; the best checkpoint was restored before testing. When few training labels were budgeted (N ≤ 1,000) we instead used 6,000 steps, warm-up 300, evaluation every 200.

#### 4.1.2 Ridge on one-hot sequence

946 sparse columns: 43 positions × 21 symbols (20 amino acids + gap) plus 43 insertion-indicator columns; ∼37.4 non-zeros per row. Regularisation chosen on the validation split from α ∈ {0.01, 0.1, 1, 10, 100}, solver `sparse_cg`. This is an additive (site-independent) model and serves as the no-epistasis reference.

#### 4.1.3 ESM-C frozen-feature head

We use the huggingface (https://huggingface.co/) repository model `biohub/ESMC-600M` (575M parameters, 36 layers, d = 1152). Each variant’s mutations are placed into the full AAV2 VP3 sequence for scoring. Hidden states from layers 18, 26 and 36 are mean-pooled over the region’s tokens only and concatenated.

For trained ESM models we use a simple MLP head, without updating the base-model’s parameters: LayerNorm → Linear(3456,512) → GELU → Dropout(0.2)→ Linear(512,1); AdamW lr 10⁻³, weight decay 0.01, cosine schedule; 4,000 steps at batch 256 for N ≤ 1,000 and 15,000 at batch 512 above; best-on-validation checkpoint.

#### 4.1.4 ESM-C zero-shot

Likelihood-based scores from the same model through a forward pass. Masked-marginal and WT-marginal scores calculated only for the 68,880 substitution-only variants, because they require a position-wise substitution at fixed length. The full-coverage alternative, unmasked pseudo-log-likelihood, is reported separately.

#### 4.1.5 PoET-2 zero-shot

OpenProteinAI/PoET-2, 400 MB checkpoint from Zenodo record 16815645, under the PoET Non-Commercial Artifacts License.

### 4.2 Evaluation splits

In the *trained on this study* rows every training and validation row comes from this study's library while the test rows come from every library. In the pooled one, train samples come from every AAV library.

### 4.3 Test splits

**Random 10%**: The target set is a 10% sample of the assayed data and its composition matches the full sample mixture. It measures interpolation.

**Fitness > +2**: train on samples which score under 2, test above it. Measures the ability to generalize beyond scores in training distribution. The hardest and most faithful challenge for therapeutic design.

**Distance ≥ 20:** train on ≤ 15 mutations, test on ≥ 20, with mutation distances 16–19 excluded entirely (79,252 variants) as a buffer so no test variant is one or two substitutions from a training variant. Measures how well models can be used for predicting diversification.

**Natural diversity:** 2,485 held-out designs from Sinai et al. These samples are all derived from importance sampling the dependoparvovirus MSA or using a VAE to generate similar sequences. Measures how well this data can predict sequences derived from evolutionary background.

**Unseen seeds:** 3,155 variants from this study’s library, forming the complete single-edit neighbourhood of three seed sequences withheld whole. Seeds 3, 9 and 14 are held out and the other sequences that are at least 9 mutations away are trained on. They are more isolated than the seeds retained for training: mean edit distance to the nearest *training* seed is 13.7, against 10.9 among the training seeds themselves (seed 14 is the most remote, at 17). They also span the library’s insertion regimes, one substitution-only, one partly carrying insertions, one heavily changed with insertions. Measures the ability to predict a targeted set of seeds, given sequences within a landscape.

The best predictive model reported is an ensemble of 5 identical models differing only in weight initialisation and data order (same data). Predictions are averaged ensemble scores.

Performance on the fitness > +2 is partly explained by the number of insertions; i.e fitness in the tail is negatively rank-correlated with insertion count (ρ = −0.277). Stratifying by insertion count and taking the n-weighted mean of the Spearman within each stratum with n ≥ 200, reduces the performance of the transformer ensemble from 0.281 to 0.176, and removes all of PoET-2’s predictive power (from 0.27 to 0.02).

## Data Availability

Experimental data for all experiments will be deposited on a public repository prior to peer-reviewed publication.

## Code Availability

The TensorFlow 1.3 API was used to implement and train original design models using the architectures described in Methods. Analysis code will be available prior to peer-reviewed publication.

## Acknowledgements

The authors would like to thank David Ding and Jeffrey Gerold for helpful discussions about this paper. The authors thank the Wyss Institute for funding.

## Author Contributions

SS, EK, GMC, LJC, AB conceived the study. EK and NJ performed in vitro experiments. DHB, AB and LJC designed, implemented and used generative ML models to generate variants with input from SS and EK. SS and EK designed the landscape exploration strategy with input from LJC and AB, selected seeds and designed the stochastic walks. SS, AG, MB, LJC analyzed the data. SS and AG wrote the analysis code and trained benchmarking models. SS, EK, and LJC wrote the paper with contributions from all authors. EK, GMC, LJC secured funding and institutional support for the project.

## Statement on the use of Large Language Models

We used Large Language Models to assist in several analyses after the experimental data was collected and parsed. All initial analysis leading to the main insights was done by the authors. LLMs were used to review and improve the analysis, the quality of plots, organize and test the code for better reproducibility, and help fix any inconsistencies in text not caught by humans. All authors take full responsibility for the entire content of the manuscript, including any segments developed with assistance from AI tools, and have reviewed and verified the results in this paper.

---

**Figure S1.**
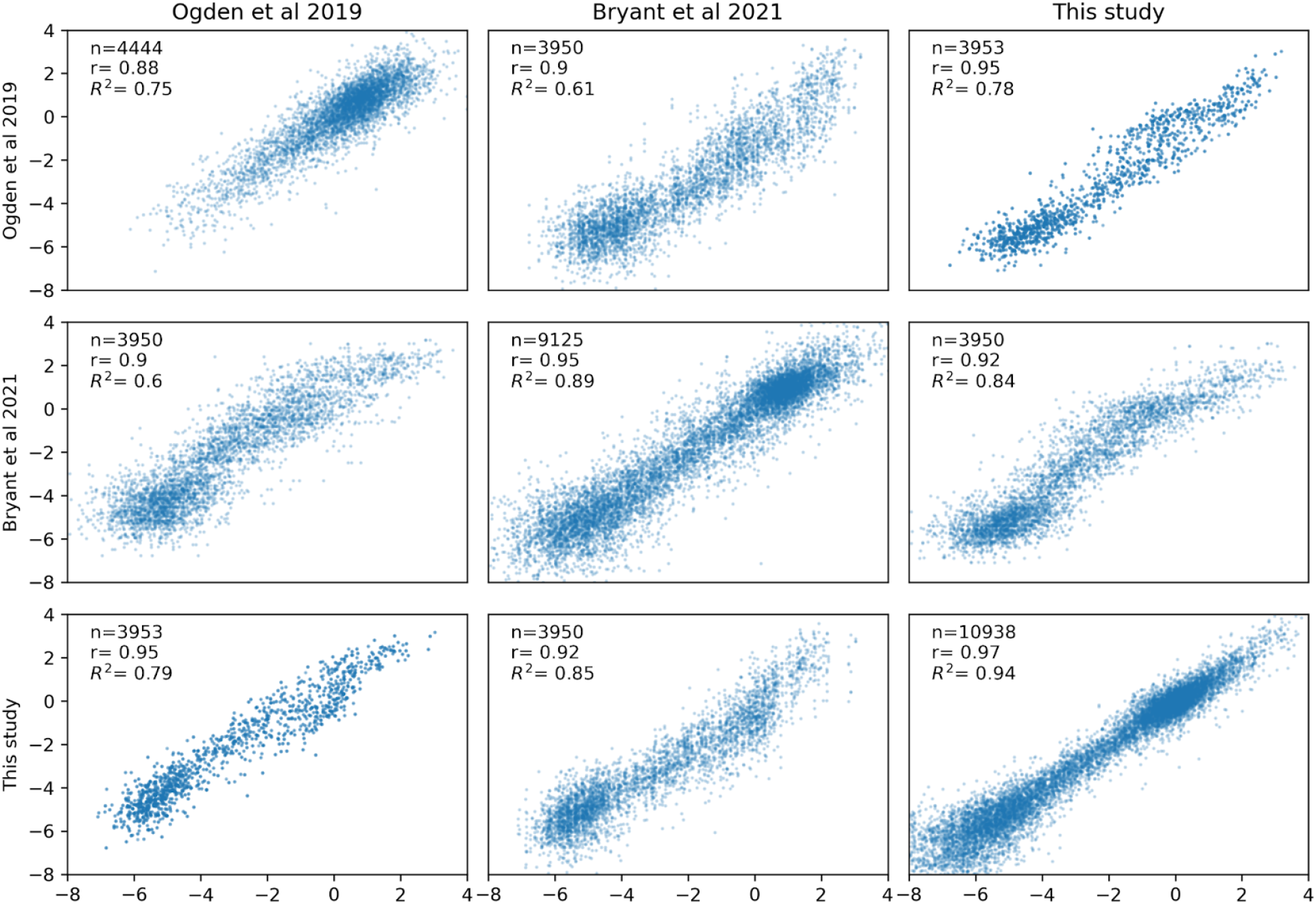
Reproducibility between capsid productivity experiments conducted in (Ogden et al. 2019), (Bryant et al. 2021) and this study. Here we compare synthetic variants at the level of nucleotide replicates synthesized separately. Because we observe high replication among the experiments in this study and those published previously we can overcome measurement noise and use quantitative traits and regression models in aggregate (in Bryant et al classification was used to avoid noise effects, we cautiously avoid analyzing single measurements in isolation in this study).

**Figure S2.**
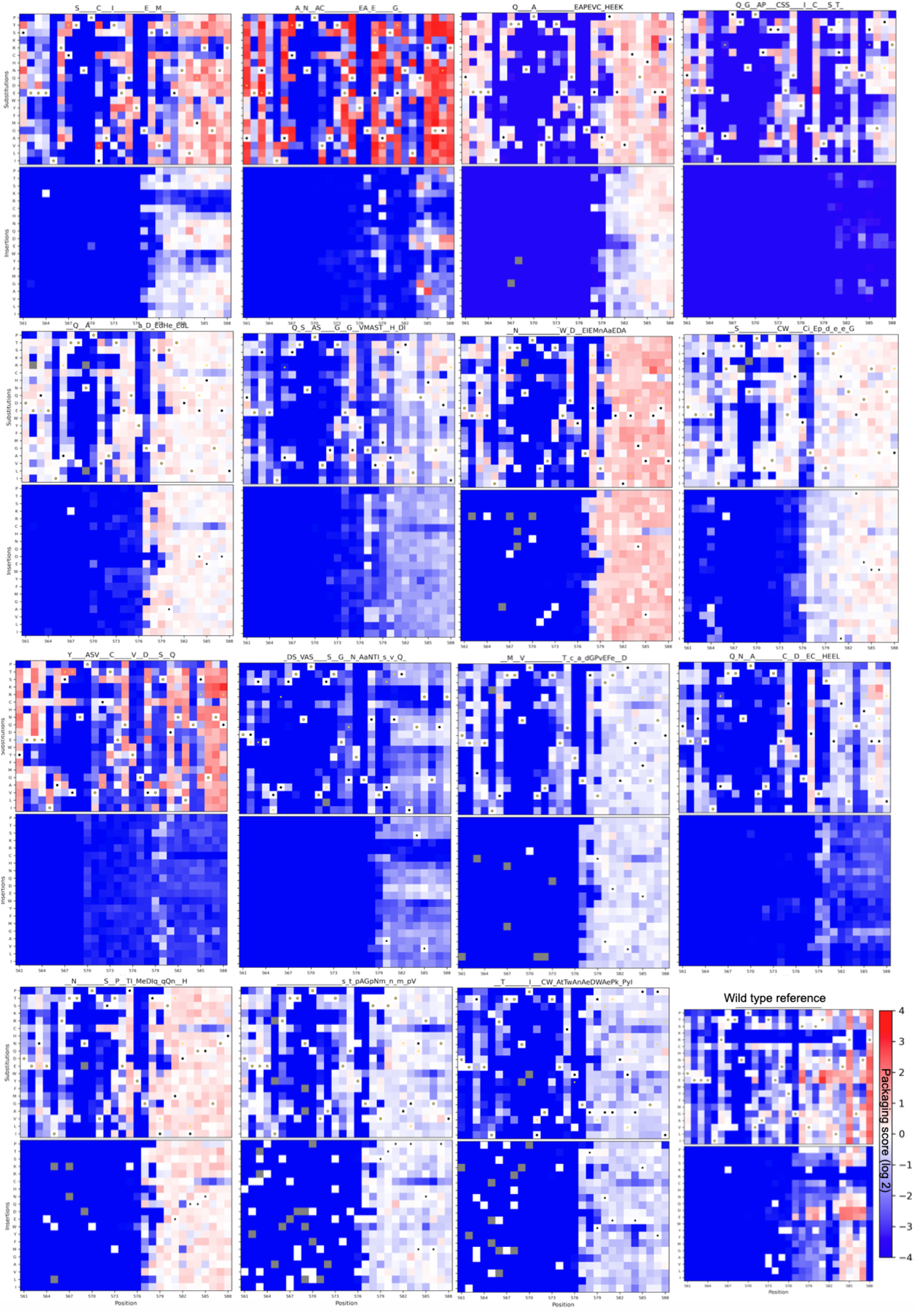
Single mutation effects around every seed. Observing single mutation effects around every seed and wild type reveals both similarity and difference in how single mutants affect each seed’s fitness (some panels are repeated from the main text).

**Figure S3.**
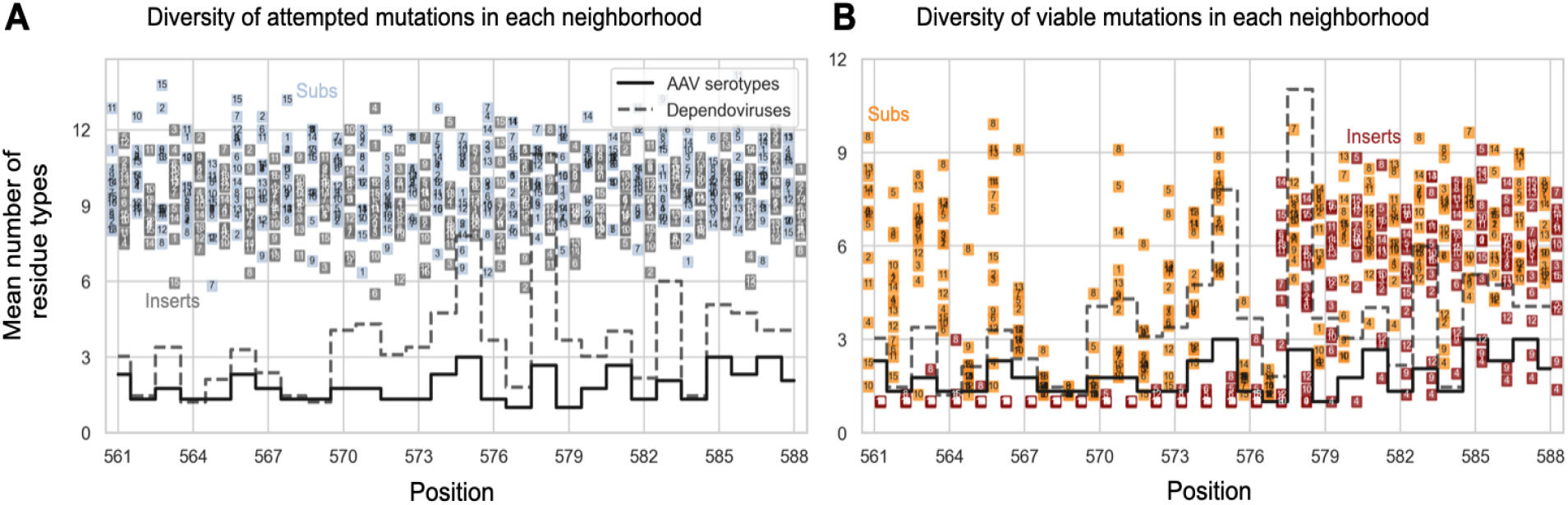
Mean number of residue types attempted vs viable. A: Mean number of residue types attempted for each seed and position (marginal distribution over single and random multi-mutants) B: Mean number of residue types that were viable. Substitutions and insertions are offset and colored separately, and each seed number is visible on the point itself.

**Figure S4.**
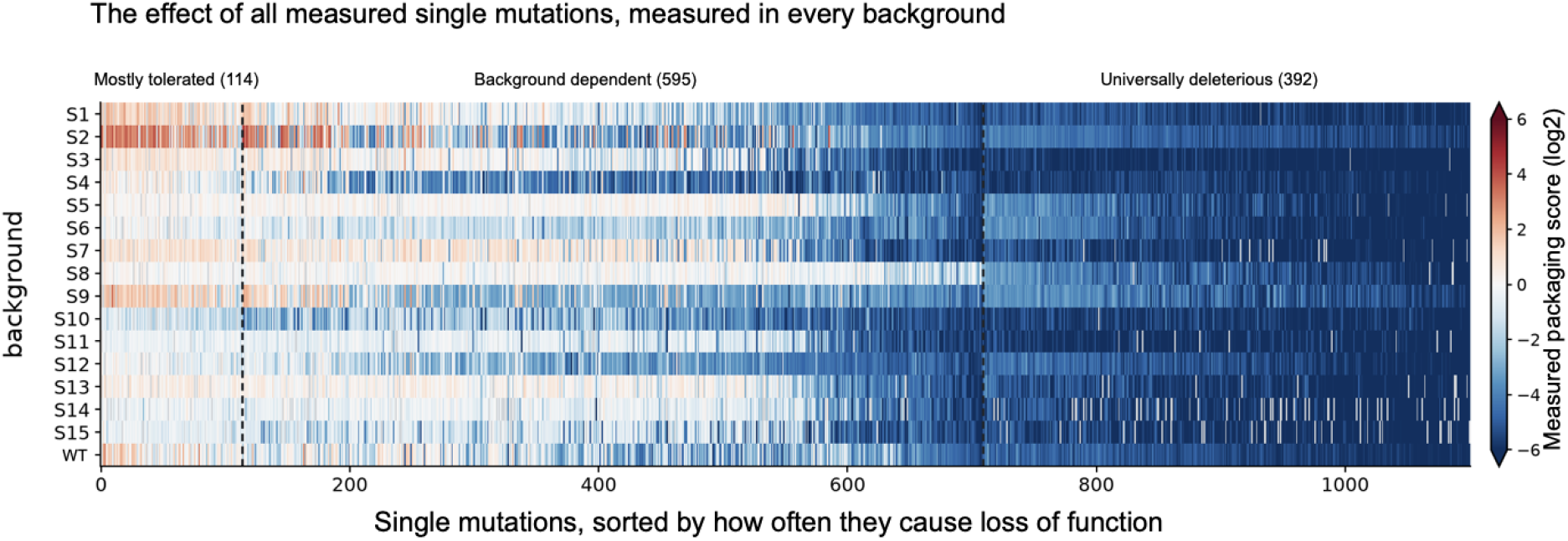
There is a set of single mutations that universally break the capsid. Single mutations around each seed and WT can be grouped into those that are mostly tolerated, those whose tolerance is highly background dependent, and those that are almost never tolerated as single mutations in the 15 + WT backgrounds. This last set is generally avoided by all other successful sampling done by other methods. Within all tested variants in this experiment, only 0.63% of those that carry one or more of these universally lethal mutations are functional (and only 0.72% with exactly one lethal mutant survive), while this number is 68.74% for variants that don’t carry any of these 392 mutations (a 109-fold change in viability).

**Figure S5.**
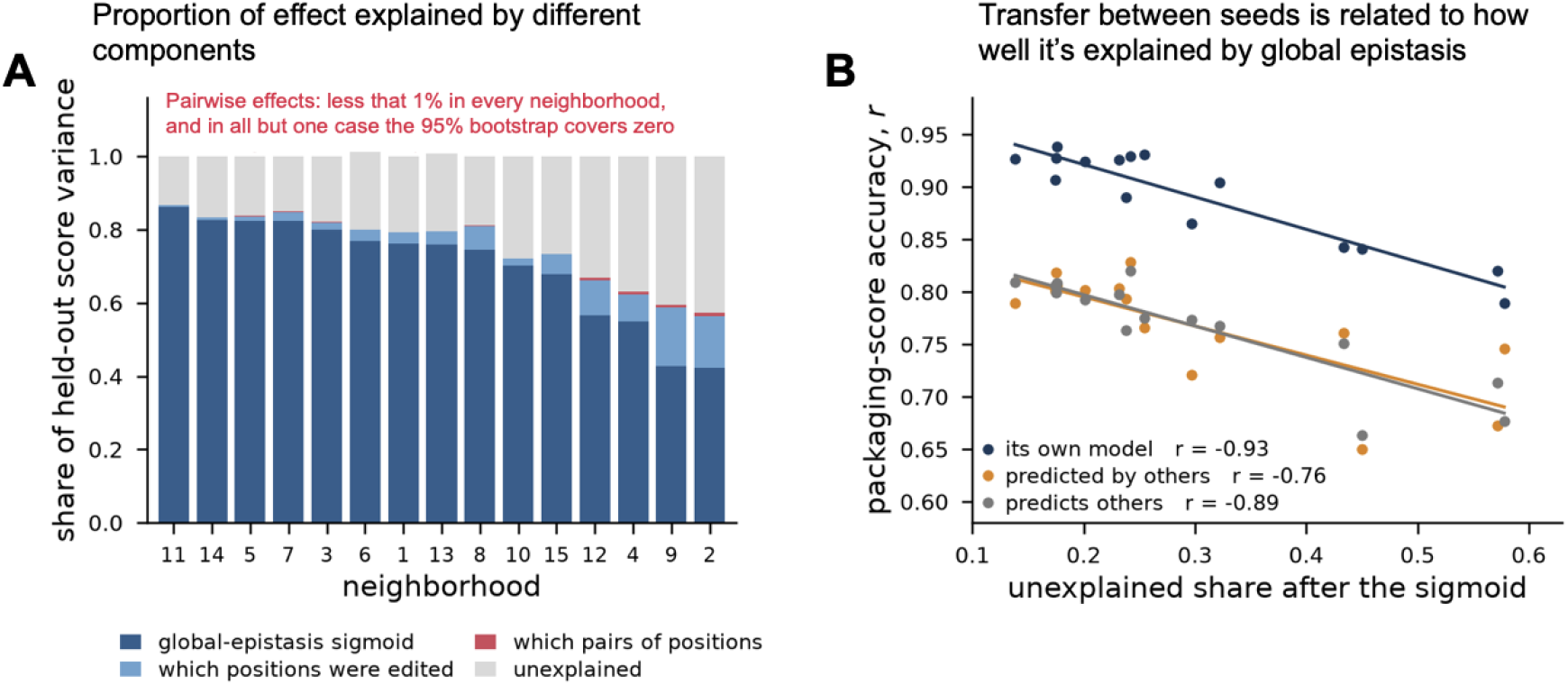
The majority of non-additive effects are explained by global epistasis. We compute the residual score remaining after we account for global epistasis, and regress on 57 positional changes plus all 1,596 distinct position pairs using ridge, with penalties chosen on that neighborhood’s validation split and evaluated on a set of ∼1000 held out multi-mutants per neighborhood. (A) we show the held out score attributable to each component, and the remaining unexplained fraction. (B) The unexplained share of a neighborhood’s score is correlated with how well that neighborhood is predicted by others. Notably, the gap between the self-model and other-models does not change depending on the fraction of effects explained by global epistasis.

**Figure S6.**
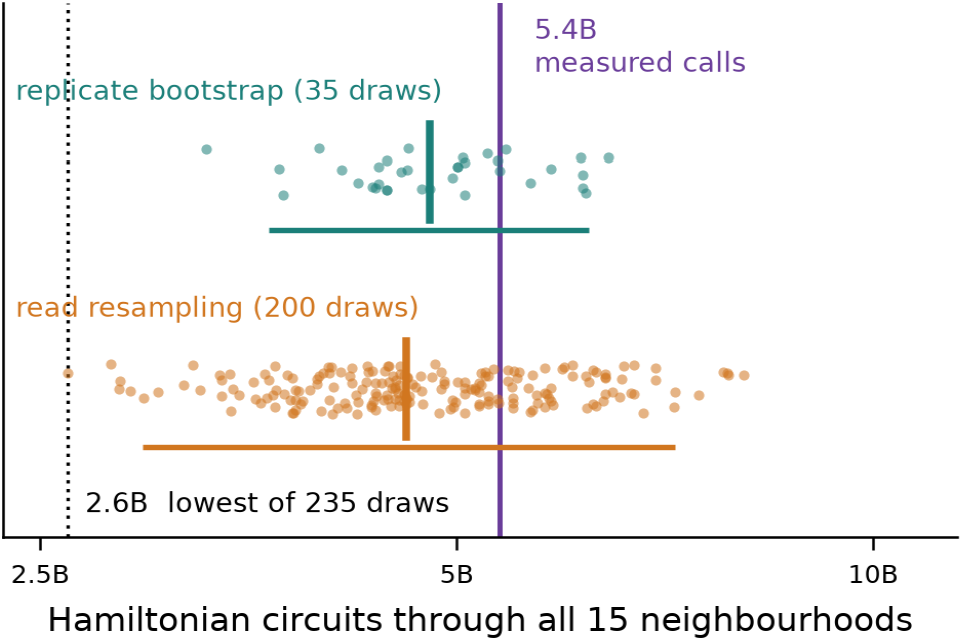
Circuit counts are robust to measurement uncertainty. To ensure the connectivity claim is robust we recompute Hamiltonian circuit counts by resampling the data through an exhaustive bootstrap over 4 sequencing replicates (35 distinct with-replacement draws, top) and Poisson resampling of plasmid and virus read counts (200 draws, bottom). Wild type score and the viability cutoff are recomputed within each sample, and each subsample was passed through the same graph construction. The purple line shows the estimate when computed from the full measurement. All 235 draws connect all 15 neighbourhoods.

**Table S1.**

| library | variants |
| --- | --- |
| Ogden et al. | 65,945 |
| Bryant et al. | 208,023 |
| **This study** | 212,558* |
| Sinai et al. | 7,290 |

\*The number of unique variants contributed to the overall pool by this study. The number of variants passing the QC filters of at least 25 plasmids (as reported in the abstract) is 232,123.

**Table S2.** Composition of train and test libraries.

| Test set | Source | train | val | test | test from this study |
| --- | --- | --- | --- | --- | --- |
| random 10% | pooled (all libraries) | 394,951 | 49,354 | 49,511 | 43.1% |
| random 10% | trained on this study | 169,872 | 21,340 | 49,511 | 43.1% |
| fitness > +2 | pooled (all libraries) | 418,779 | 46,590 | 28,447 | 3.9% |
| fitness > +2 | trained on this study | 190,216 | 21,241 | 28,447 | 3.9% |
| distance 20–28 | pooled (all libraries) | 295,248 | 32,789 | 86,527 | 28.4% |
| distance 20–28 | trained on this study | 129,775 | 14,468 | 86,527 | 28.4% |
| natural diversity | pooled (all libraries) | 442,203 | 49,128 | 2,485 | 0.0% |
| natural diversity | trained on this study | 191,218 | 21,340 | 2,485 | 0.0% |
| unseen seeds | pooled (all libraries) | 418,267 | 46,494 | 3,155 | 100.0% |
| unseen seeds | trained on this study | 166,283 | 18,615 | 3,155 | 100.0% |

**Table S3.**
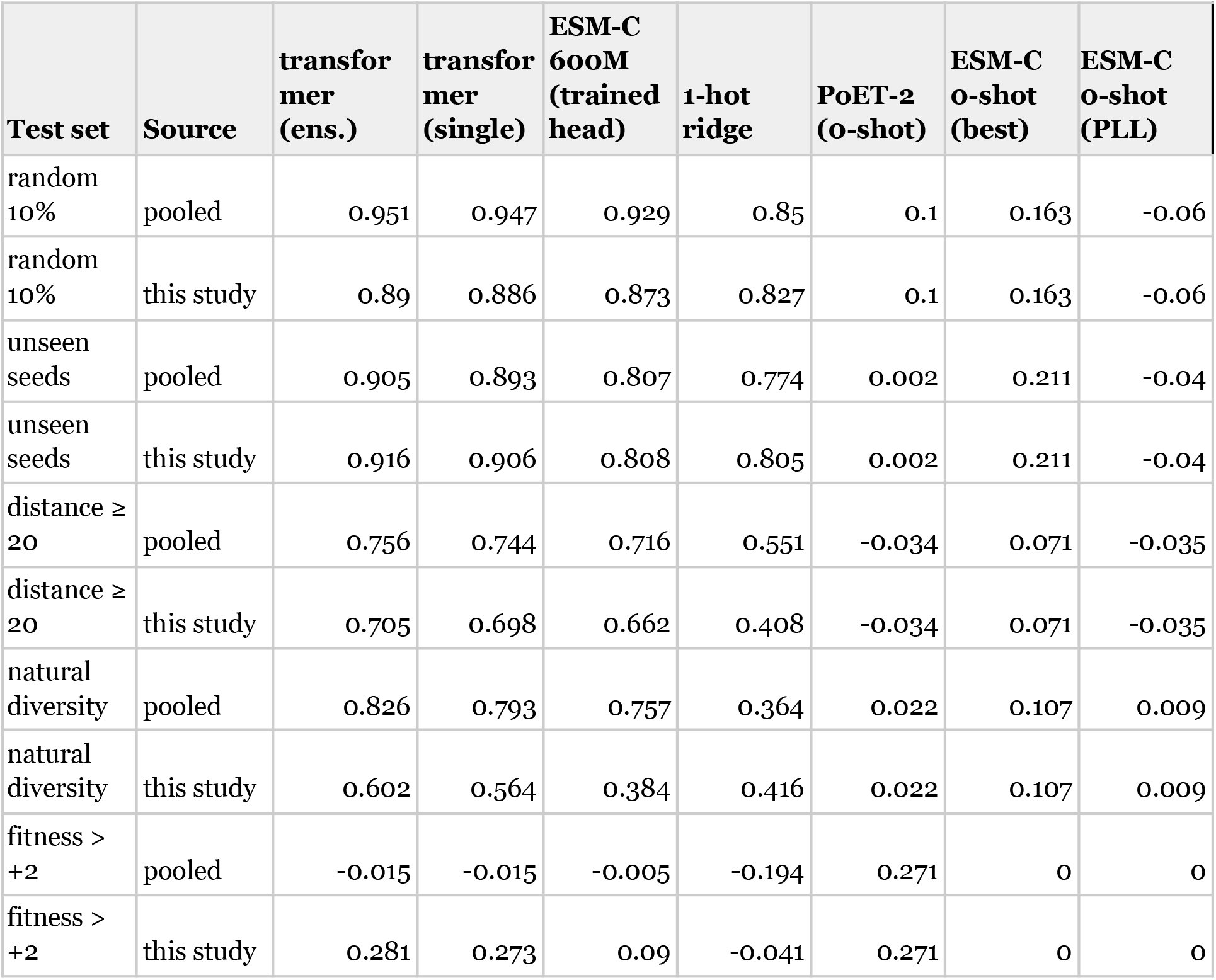
Performance of models on each test set split.

## References

1. Adachi, Kei, Tatsuji Enoki, Yasuhiro Kawano, Michael Veraz, and Hiroyuki Nakai. 2014. “Drawing a High-Resolution Functional Map of Adeno-Associated Virus Capsid by Massively Parallel Sequencing.” Nature Communications 5: 3075. 10.1038/ncomms4075.

2. Avizemer, Ziv, Carlos Martí-Gómez, Shlomo Yakir Hoch, David M. McCandlish, and Sarel J. Fleishman. 2025. “Evolutionary Paths That Link Orthogonal Pairs of Binding Proteins.” Cell Systems 16 (5): 101262. 10.1016/j.cels.2025.101262.

3. Bakerlee, Christopher W., Alex N. Nguyen Ba, Yekaterina Shulgina, Jose I. Rojas Echenique, and Michael M. Desai. 2022. “Idiosyncratic Epistasis Leads to Global Fitness-Correlated Trends.” Science 376 (6593): 630–35. 10.1126/science.abm4774.

4. Bank, Claudia, Sebastian Matuszewski, Ryan T. Hietpas, and Jeffrey D. Jensen. 2016. “On the (Un)Predictability of a Large Intragenic Fitness Landscape.” Proceedings of the National Academy of Sciences 113 (49): 14085–90. 10.1073/pnas.1612676113.

5. Biswas, Surojit, Grigory Khimulya, Ethan C. Alley, Kevin M. Esvelt, and George M. Church. 2021. “Low-N Protein Engineering with Data-Efficient Deep Learning.” Nature Methods 18 (4): 389–96. 10.1038/s41592-021-01100-y.

6. Brookes, David H., Amirali Aghazadeh, and Jennifer Listgarten. 2022. “On the Sparsity of Fitness Functions and Implications for Learning.” Proceedings of the National Academy of Sciences of the United States of America 119 (1). 10.1073/pnas.2109649118.

7. Brookes, David, Jakub Otwinowski, and Sam Sinai. 2024. “Contrastive Losses as Generalized Models of Global Epistasis.” Advances in Neural Information Processing Systems 37, 93374–405. 10.52202/079017-2962.

8. Bryant, Drew H., Ali Bashir, Sam Sinai, et al. 2021. “Deep Diversification of an AAV Capsid Protein by Machine Learning.” Nature Biotechnology 39 (6): 691–96. 10.1038/s41587-020-00793-4.

9. Candido, Salvatore, Thomas Hayes, Alexander Derry, et al. 2026. “Language Modeling Materializes a World Model of Protein Biology.” In *bioRxiv*. bioRxiv, June 4. 10.64898/2026.06.03.729735.

10. Ding, David, Anna G. Green, Boyuan Wang, et al. 2022. “Co-Evolution of Interacting Proteins through Non-Contacting and Non-Specific Mutations.” Nature Ecology & Evolution 6 (5): 590–603. 10.1038/s41559-022-01688-0.

11. Ding, David, Ada Y. Shaw, Sam Sinai, et al. 2024. “Protein Design Using Structure-Based Residue Preferences.” Nature Communications 15 (1): 1639. 10.1038/s41467-024-45621-4.

12. Ding, Frances, and Jacob Steinhardt. 2024. “Protein Language Models Are Biased by Unequal Sequence Sampling across the Tree of Life.” In bioRxiv. 10.1101/2024.03.07.584001.

13. Diplock, Nathan, Thomas Desautels, Riyaz Haque, et al. 2026. “Unobserved Sequence Space Has Many Functional Proteins.” In bioRxiv. bioRxiv, August 26. 10.64898/2026.08.24.746828.

14. Freschlin, Chase R., Sarah A. Fahlberg, Pete Heinzelman, and Philip A. Romero. 2024. “Neural Network Extrapolation to Distant Regions of the Protein Fitness Landscape.” Nature Communications 15 (1): 6405. 10.1038/s41467-024-50712-3.

15. Gelman, Sam, Sarah A. Fahlberg, Pete Heinzelman, Philip A. Romero, and Anthony Gitter. 2021. “Neural Networks to Learn Protein Sequence–Function Relationships from Deep Mutational Scanning Data.” Proceedings of the National Academy of Sciences 118 (48): e2104878118. 10.1073/pnas.2104878118.

16. Gonzalez Somermeyer, Louisa, Aubin Fleiss, Alexander S. Mishin, et al. 2022. “Heterogeneity of the GFP Fitness Landscape and Data-Driven Protein Design.” eLife 11 (May). 10.7554/eLife.75842.

17. Greenbury, Sam F., Ard A. Louis, and Sebastian E. Ahnert. 2022. “The Structure of Genotype-Phenotype Maps Makes Fitness Landscapes Navigable.” Nature Ecology & Evolution 6 (11): 1742–52. 10.1038/s41559-022-01867-z.

18. Hopf, Thomas A., John B. Ingraham, Frank J. Poelwijk, et al. 2017. “Mutation Effects Predicted from Sequence Co-Variation.” Nature Biotechnology 35 (2): 128–35. 10.1038/nbt.3769.

19. Hsu, Chloe, Hunter Nisonoff, Clara Fannjiang, and Jennifer Listgarten. 2022. “Learning Protein Fitness Models from Evolutionary and Assay-Labeled Data.” Nature Biotechnology 40 (7): 1114–22. 10.1038/s41587-021-01146-5.

20. Huang, Qin, Ken Y. Chan, Jason Wu, et al. 2024. “An AAV Capsid Reprogrammed to Bind Human Transferrin Receptor Mediates Brain-Wide Gene Delivery.” Science (New York, N.Y.) 384 (6701): 1220–27. 10.1126/science.adm8386.

21. Jiang, Kaiyi, Zhaoqing Yan, Matteo Di Bernardo, et al. 2024. “Rapid in Silico Directed Evolution by a Protein Language Model with EVOLVEpro.” Science (New York, N.Y.), November 21, eadr6006. 10.1126/science.adr6006.

22. Kauffman, S. A., and E. D. Weinberger. 1989. “The NK Model of Rugged Fitness Landscapes and Its Application to Maturation of the Immune Response.” Journal of Theoretical Biology 141 (2): 211–45. 10.1016/s0022-5193(89)80019-0.

23. Kinney, Justin B. 2026. “Mount Fuji’s Stubby Peak: The Genotypic Density of Additive Landscapes near Maximal Fitness.” Genetics, no. iyag133 (June). 10.1093/genetics/iyag133.

24. Klug, Alexander, Su-Chan Park, and Joachim Krug. 2019. “Recombination and Mutational Robustness in Neutral Fitness Landscapes.” PLoS Computational Biology 15 (8): e1006884. 10.1371/journal.pcbi.1006884.

25. Madani, Ali, Ben Krause, Eric R. Greene, et al. 2023. “Large Language Models Generate Functional Protein Sequences across Diverse Families.” Nature Biotechnology 41 (8): 1099–106. 10.1038/s41587-022-01618-2.

26. Nimwegen, E. van, J. P. Crutchfield, and M. Huynen. 1999. “Neutral Evolution of Mutational Robustness.” Proceedings of the National Academy of Sciences of the United States of America 96 (17): 9716–20. 10.1073/pnas.96.17.9716.

27. Ogden, Pierce J., Eric D. Kelsic, Sam Sinai, and George M. Church. 2019. “Comprehensive AAV Capsid Fitness Landscape Reveals a Viral Gene and Enables Machine-Guided Design.” Science 366 (6469): 1139–43. 10.1126/science.aaw2900.

28. Oros, Daniel, and Joachim Krug. 2026. “Random Fitness Landscapes Are Highly Navigable.” In *bioRxiv*. bioRxiv, May 26. 10.64898/2026.05.22.727193.

29. Otwinowski, Jakub, David M. McCandlish, and Joshua B. Plotkin. 2018. “Inferring the Shape of Global Epistasis.” Proceedings of the National Academy of Sciences of the United States of America 115 (32): E7550–58. 10.1073/pnas.1804015115.

30. Papkou, Andrei, Lucia Garcia-Pastor, José Antonio Escudero, and Andreas Wagner. 2023. “A Rugged yet Easily Navigable Fitness Landscape.” Science 382 (6673): eadh3860. 10.1126/science.adh3860.

31. Park, Yeonwoo, Brian P. H. Metzger, and Joseph W. Thornton. 2024. “The Simplicity of Protein Sequence-Function Relationships.” Nature Communications 15 (1): 7953. 10.1038/s41467-024-51895-5.

32. Poelwijk, Frank J., Michael Socolich, and Rama Ranganathan. 2019. “Learning the Pattern of Epistasis Linking Genotype and Phenotype in a Protein.” Nature Communications 10 (1): 4213. 10.1038/s41467-019-12130-8.

33. Puszkarska, Anna M., Bruck Taddese, Jefferson Revell, et al. 2024. “Machine Learning Designs New GCGR/GLP-1R Dual Agonists with Enhanced Biological Potency.” Nature Chemistry 16 (9): 1436–44. 10.1038/s41557-024-01532-x.

34. Reddy, Gautam, and Michael M. Desai. 2021. “Global Epistasis Emerges from a Generic Model of a Complex Trait.” eLife 10 (March): e64740. 10.7554/eLife.64740.

35. Romero, Philip A., Andreas Krause, and Frances H. Arnold. 2013. “Navigating the Protein Fitness Landscape with Gaussian Processes.” Proceedings of the National Academy of Sciences of the United States of America 110 (3): E193–201. 10.1073/pnas.1215251110.

36. Rotrattanadumrong, Rachapun, and Yohei Yokobayashi. 2022. “Experimental Exploration of a Ribozyme Neutral Network Using Evolutionary Algorithm and Deep Learning.” Nature Communications 13 (1): 4847. 10.1038/s41467-022-32538-z.

37. Russ, William P., Matteo Figliuzzi, Christian Stocker, et al. 2020. “An Evolution-Based Model for Designing Chorismate Mutase Enzymes.” *Science (New York*, N.Y.) 369 (6502): 440–45. 10.1126/science.aba3304.

38. Sailer, Zachary R., and Michael J. Harms. 2017. “Detecting High-Order Epistasis in Nonlinear Genotype-Phenotype Maps.” Genetics 205 (3): 1079–88. 10.1534/genetics.116.195214.

39. Sarkisyan, Karen S., Dmitry A. Bolotin, Margarita V. Meer, et al. 2016. “Local Fitness Landscape of the Green Fluorescent Protein.” Nature 533 (7603): 397–401. 10.1038/nature17995.

40. Schuster, Peter, and Peter F. Stadler. 2023. “Virus Evolution on Fitness Landscapes.” Current Topics in Microbiology and Immunology 439: 1–94. 10.1007/978-3-031-15640-3_1.

41. Sinai, Sam, Nina Jain, George M. Church, and Eric D. Kelsic. 2021. “Generative AAV Capsid Diversification by Latent Interpolation.” In bioRxiv. 10.1101/2021.04.16.440236.

42. Tabebordbar, Mohammadsharif, Kim A. Lagerborg, Alexandra Stanton, et al. 2021. “Directed Evolution of a Family of AAV Capsid Variants Enabling Potent Muscle-Directed Gene Delivery across Species.” Cell 184 (19): 4919–4938.e22. 10.1016/j.cell.2021.08.028.

43. Truong, Timothy Fei, Jr, and Tristan Bepler. 2025. “Understanding Protein Function with a Multimodal Retrieval-Augmented Foundation Model.” In arXiv [q-Bio.QM]. August 5. arXiv. 10.48550/arXiv.2508.04724.

44. Wagner, Andreas. 2008. “Neutralism and Selectionism: A Network-Based Reconciliation.” Nature Reviews. Genetics 9 (12): 965–74. 10.1038/nrg2473.

45. Weinreich, Daniel M., Nigel F. Delaney, Mark A. Depristo, and Daniel L. Hartl. 2006. “Darwinian Evolution Can Follow Only Very Few Mutational Paths to Fitter Proteins.” Science 312 (5770): 111–14. 10.1126/science.1123539.

46. Weinreich, Daniel M., Yinghong Lan, C. Scott Wylie, and Robert B. Heckendorn. 2013. “Should Evolutionary Geneticists Worry about Higher-Order Epistasis?” Current Opinion in Genetics & Development 23 (6): 700–707. 10.1016/j.gde.2013.10.007.

47. Wright, Sewall, and Others. 1932. The Roles of Mutation, Inbreeding, Crossbreeding, and Selection in Evolution. http://www.esp.org/books/6th-congress/facsimile/contents/6th-cong-p356-wright.pdf.

48. Wu, Zachary, S. B. Jennifer Kan, Russell D. Lewis, Bruce J. Wittmann, and Frances H. Arnold. 2019. “Machine Learning-Assisted Directed Protein Evolution with Combinatorial Libraries.” Proceedings of the National Academy of Sciences of the United States of America 116 (18): 8852–58. 10.1073/pnas.1901979116.

49. Zhou, Juannan, and David M. McCandlish. 2020. “Minimum Epistasis Interpolation for Sequence-Function Relationships.” Nature Communications 11 (1): 1782. 10.1038/s41467-020-15512-5.

50. Zhou, Juannan, Mandy S. Wong, Wei-Chia Chen, Adrian R. Krainer, Justin B. Kinney, and David M. McCandlish. 2022. “Higher-Order Epistasis and Phenotypic Prediction.” Proceedings of the National Academy of Sciences of the United States of America 119 (39): e2204233119. 10.1073/pnas.2204233119.

51. Zhu, Danqing, David H. Brookes, Akosua Busia, et al. 2024. “Optimal Trade-off Control in Machine Learning–Based Library Design, with Application to Adeno-Associated Virus (AAV) for Gene Therapy.” Science Advances 10 (4): eadj3786. 10.1126/sciadv.adj3786.

